# Eco-evolutionary dynamics in competitive systems: Rescue and murder

**DOI:** 10.64898/2026.01.14.699340

**Authors:** Simon Leoz, Frithjof Lutscher, Korinna T. Allhoff, Lynn Govaert

## Abstract

Rapid evolution in response to changing environments can have beneficial (e.g. evolutionary rescue) or detrimental (e.g. evolutionary suicide) outcomes for the survival of one species. Responses of multi-species systems are even harder to predict, but important to consider. We simulate eco-evolutionary dynamics in one- and two-species systems, with two traits per species: physiological performance (associated with abiotic pressure) and defense against interference competition (associated with biotic pressure). In the single-species system, evolution is consistently beneficial, enabling evolutionary rescue. In contrast, in the two-species system, evolution can be simultaneously beneficial and detrimental, because rescue of one species can lead to evolutionary murder of the other. Notably, the pattern of rescue and murder was largely independent of different ecological and evolutionary scenarios, indicating that evolutionary murder might be common in competitive systems. Our study enables exploration of eco-evolutionary dynamics in more complex biotic settings, extending understanding of species responses to abiotic and biotic changes.

## 1 Introduction

Global change is severely impacting ecosystems and threatening ecosystem functions (Hooper et al. 2012; Bernhardt et al. 2017; Pascual et al. 2022). Understanding how ecosystems respond to these changing conditions will give important insights into maintaining biodiversity and well-functioning ecosystems (Sandifer et al. 2015; Oliveira et al. 2022). Historically, this challenge has been addressed from a purely ecological perspective, which assumes that ecological and evolutionary processes act on separate timescales. In recent decades, however, increasing evidence has shown that rapid evolutionary responses can occur within just a few generations (Hairston et al. 2005; Koch et al. 2014), especially under fast-changing anthropogenic pressures and disturbances (Hendry et al. 2008; Hoffmann & Sgrò 2011), potentially resulting in eco-evolutionary feedbacks (Post & Palkovacs 2009; Schoener 2011; Hendry 2017). Rapid evolution has been shown to affect ecological dynamics at the population (Yoshida et al. 2003), community (Pantel et al. 2015), and ecosystem (Bassar et al. 2012) level. However, whether rapid evolution and the resulting eco-evolutionary dynamics foster or hinder species persistence in changing environments remains to be fully explored (Pelletier et al. 2009).

Rapid evolution can have beneficial effects for species persistence, e.g., when a population avoids extinction due to severe external stress through adaptation by natural selection, a process called evolutionary rescue (Gomulkiewicz & Holt 1995; Bell & Gonzalez 2009; Carlson et al. 2014; Bell 2017). Empirical examples of such evolutionary rescue range from adaptation to water acidification for phytoplankton species to evolution of vertebrate species in response to pesticides and viruses (Bell 2013; Vander Wal et al. 2013; Bell 2017). However, several theoretical studies have also shown that evolution can negatively affect population survival, potentially leading to evolutionary suicide, when a viable population evolves traits that ultimately prevent its persistence (Parvinen 2005; Ferriere & Legendre 2013). Together, these concepts highlight how evolution can produce both beneficial and detrimental outcomes in altered environments.

In multi-species settings, the question of whether evolution promotes or hinders species persistence is even more complex. Evolutionary rescue can be both direct (as described earlier) or indirect. For example, a non-evolving predator can be rescued from extinction through evolution of its prey, when a defence-growth trade-off in the prey population exists (Yamamichi & Miner 2015). In small food web modules, indirect rescue of others can even occur more frequently than self-rescue (van Velzen 2023). However, the evolution of one species can also drive another species to extinction, resulting in evolutionary murder (Parvinen 2005). For instance, plants that experience declining pollinator abundances might invest less into attracting those pollinators and more into selfing or vegetative reproduction, which in turn further threatens the already declining pollinator population (Weinbach et al. 2022; Weyerer et al. 2023). Evolutionary murder can also occur between trophic levels, such as in the case of systems where the prey’s adaptation affects the persistence of a predator subject to experiencing external disturbance (Shang et al. 2024). We hypothesize that similar scenarios of evolutionary murder could occur in competitive systems. For example, when a competing species can rapidly adapt to changes in the abiotic environment, its resulting high abundances can increase competitive pressure on the focal species. This increased competitive pressure can then accelerate the decline or even extinction of the competitor, especially if their adaptation to the abiotic changes lags behind.

Given that all species are embedded within communities, it is important to consider species interactions when studying species responses to environmental change (De Meester et al. 2019; Loeuille 2019). Indeed, previous studies have shown that biotic interactions can induce rapid evolutionary changes (Yoshida et al. 2003) as well as alter evolutionary responses to abiotic environmental change (Lawrence et al. 2012; Govaert et al. 2021; Grainger et al. 2021). Within a community, species experience both biotic and abiotic selective pressures. Trait evolution in response to abiotic or biotic change, or both, often occurs for multiple traits simultaneously (Stoks et al. 2016), and the speed at which such multi-variate trait evolution can occur depends on the underlying genetic variation and constraints of these traits (Lande 1979; Schluter 1996). Traits differ in their genetic variation (Blows & Hoffmann 2005), and evolutionary changes in one trait can aid or hamper evolutionary changes in other traits (Hine et al. 2014). For example, in competitive systems, where competition can promote trait divergence, genetic covariance can affect evolutionary rescue by facilitating character displacement (Morita & Yamamichi 2021). Yet, how such genetic trade-offs determine eco-evolutionary outcomes, including evolutionary murder, remains an open question.

In this study, we use a two-species Lotka-Volterra competition model to investigate whether the effects of evolution on species persistence are beneficial or detrimental, and under what conditions eco-evolutionary feedbacks speed up or slow down population decline in a changing environment. Specifically, each species in our model has two traits that can evolve: one linked to its physiological performance in the abiotic environment and one linked to its defence against competitive pressure. The abiotic environment deteriorates progressively so that, in the absence of evolution, both species eventually become extinct. The exact time point of extinction, however, depends on how quickly species adapt to the changing abiotic and biotic conditions. The extent to which such adaptation can occur, in turn, depends on both evolutionary and ecological constraints. In the absence of interspecific competition, our model simplifies into a single-species system. We first demonstrate that our model results are then consistent with predictions from evolutionary rescue theory (Gomulkiewicz & Holt 1995; Bell & Gonzalez 2009; Bell 2017). The novelty of our study lies in analyzing the full 2-species competitive system, where we show how evolution can lead to both evolutionary rescue and murder. Our model provides general insights into how the interplay between biotic and abiotic selection pressures shapes species density and trait trajectories and determines whether eco-evolutionary interactions can accelerate or prevent species extinction under environmental change. By tracking extinction time as an ecological outcome alongside trait evolution, our framework provides quantities that are accessible in various empirical systems.

## 2 Model and methods

Our model describes the ecological (population density) and evolutionary (trait) dynamics of two species interacting via interference competition, which can for example occur when individuals collide or block access to resources (Holdridge et al. 2016). Such model systems include, for example, rodents like gerbils (Ziv et al. 1993), flycatchers (Qvarnström et al. 2009) or freshwater ciliates (Jiang & Morin 2007; Govaert & Klauschies 2025). The species are experiencing abiotic selection pressure from a changing abiotic environment (e.g., increasing salinity levels) and biotic selection pressure from intraspecific and interspecific competitive interactions. Species can evolve their physiological optimum and defence against competition. We implement an evolutionary trade-off, reflecting underlying genetic trait constraints (Blows & Hoffmann 2005; Hine et al. 2014), as well as an ecological trade-off, representing growth-defence trade-off theory (Ehrlich et al. 2020). Model variables and parameters are summarized in Table 1. Simulations were performed in Python (version 3.11).

**Table 1:** Summary of state variables (top) and model parameters (bottom), their description, units and (initial) values. We indicate species-specific parameters with subscripts *X* and *Y*, respectively.

| Symbol | Name | Unit | Value |
| --- | --- | --- | --- |
| $x_i(t), y_j(t)$ | population density of type $i$ of species $X$, type $j$ of species $Y$ at time $t$ | $\#^* \cdot L^{-1}$ | 0.05 |
| $c(t)$ | state of the environment at time $t$ | [1] | 1 |
| $c_{X,i}, c_{Y,j}$ | physiological optimum trait | [1] | 1 |
| $a_{X,i}, a_{Y,j}$ | sensitivity to competition | $L \cdot \#^{-1}$ | 1 |
| $d_{X,i}, d_{Y,j} (d_{\max})$ | defence against competition (maximum) | $\# \cdot L^{-1}$ | 0.8 (1) |
| $r_{X,i}, r_{Y,j} (r_{\max})$ | intrinsic growth rate (maximum) | $[t]^{-1}$ | determined by Equation (3) (1) |
| $b_{XY} (b_{YX})$ | ratio between sensitivity to intra- and interspecific competitive pressure experienced by species $X$ ($Y$) | [1] | 0.8 |
| $p$ | initial mutant density / extinction threshold | $\# \cdot L^{-1}$ | $1 \cdot 10^{-3}$ |
| $\sigma_{c_X}, \sigma_{c_Y}$ | mutation spread of physiological optimum | [1] | $[0, \sigma_{\max,c}]$ |
| $\sigma_{d_X}, \sigma_{d_Y}$ | mutation spread of defence against competition | $\# \cdot L^{-1}$ | $[0, \sigma_{\max,d}]$ |
| $\sigma_{\max,c}$ | maximum mutation spread for the physiological optimum | [1] | 0.03 |
| $\sigma_{\max,d}$ | maximum mutation spread for the defence against competition | $\# \cdot L^{-1}$ | 0.03 |
| $\sigma_{w,X}, \sigma_{w,Y}$ | spread of physiological performance | [1] | 0.85 |
| $\mu_X, \mu_Y$ | per-capita mutation rate | $[t]^{-1}$ | $5 \cdot 10^{-4}$ |
| $s_{\text{eco}} (s_{\text{evo}})$ | ecological (evolutionary) trade-off shape | [1] | $\{0.5, 1, 2\}$ |
| $\tau$ | rate of abiotic environmental change | $[t]^{-1}$ | $2 \cdot 10^{-6}$ |
| $t_f$ | maximum simulation runtime | $[t]$ | $5 \cdot 10^6$ |

\* $\#$ denotes the number of individuals.

### 2.1 Ecological dynamics

For the ecological dynamics, we assume interference competition and use a Lotka-Volterra model. We denote the two competing species by *X* and *Y* and their respective densities at time *t* by *x*(*t*) and *y*(*t*) (see Table 1).

The dynamics of species *X* and *Y* are described by the differential equations:

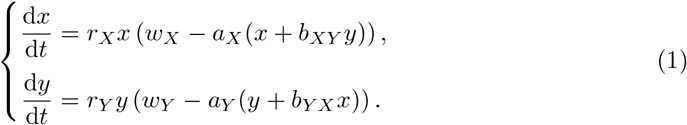

The species’ intrinsic growth rates (*r_X_, r_Y_*) are the maximum per capita growth rates under optimal conditions, while the terms in parentheses account for growth limitation due to physiological performance (abiotic environment) and competition (biotic environment).

The physiological performance of species *X* depends on the difference between the state of the environment, *c* = *c*(*t*) (e.g., salinity level), and the physiological optimum of the species, *c_X_*. It is expressed by the *physiological performance function*, *w_X_*, which we model as a Gaussian with spread σ*_w,X_* (Little & Seebacher 2021; Liu et al. 2021):

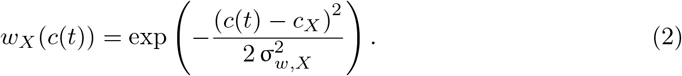

This function is positive, reaches a maximum of 1 when the physiological optimum (*c_X_*) matches the state of the environment (*c*(*t*)), and decreases when the distance between *c_X_* and *c*(*t*) increases. The parameter σ*_w,X_* corresponds to the spread of the function, with smaller values representing a narrower performance curve, and hence a larger cost when the species’ trait does not match the state of the abiotic environment. The considerations for *w_Y_* are analogous.

The second term in parentheses in Equation (1) describes intra- and interspecific competition. This term depends on a population’s sensitivity to competition (*a_X_, a_Y_*) and the ratio between intra- and interspecific competitive pressure (*b_XY_*, *b_Y X_*). We assume that *b_XY_* and *b_Y X_* are less than 1, reflecting the assumption that niche differentiation or behavioral difference reduce interspecific competition relative to intraspecific competition (MacArthur & Wilson 1967; Tilman 1982). In the following, we interpret *d_X_* = *a^−^*^1^ as the defence of species *X* against competition, and similarly for *d_Y_* = *a^−^*^1^.

Following growth-defence trade-off theory (Ehrlich et al. 2020), we implement a negative ecological trade-off between the intrinsic growth rate *r_X_* and defence *d_X_*, so that high values of *d_X_* imply low values of *r_X_* and vice versa. We let *r*_max_ and *d*_max_ be the maximal values of those parameters and express the trade-off as

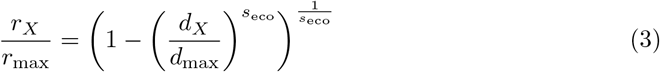

where *s*_eco_ *>* 0 determines the shape of the trade-off (Figure 1). The relation is convex, concave, or linear when *s*_eco_ *<* 1, *s*_eco_ *>* 1, or *s*_eco_ = 1, respectively. The formulation for species *Y* is analogous. Note that by affecting intrinsic growth rates, this trade-off also affects the speed of ecological dynamics.

**Figure 1:**
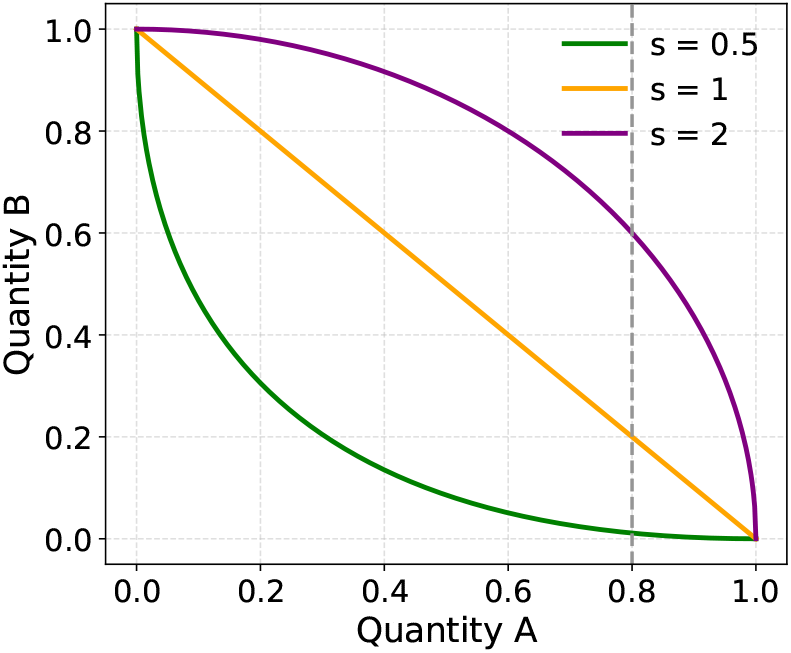
Visualization of the trade-off functions from Equations (3) and (4). For the ecological trade-off, quantity A denotes *d/d*_max_ and quantity B denotes *r/r*_max_. For the evolutionary trade-off, quantity A denotes σ*_c_/*σ_max*,c*_ and quantity B denotes σ*_d_/*σ_max*,d*_. The dashed line indicates the different initial values of *r/r*_max_ for the fixed initial value *d/d*_max_ = 0.8, as used in Figures 2-5, depending on the ecological trade-off shape.

### 2.2 Evolutionary dynamics

We model the evolution of the two traits (physiological optimum, *c_X_, c_Y_*, and defence against competition, *d_X_, d_Y_*) via recurrent mutation events. Our approach is inspired by Adaptive Dynamics (Geritz et al. 1998; Brännström et al. 2013), but we do not assume a strict separation of ecological and evolutionary timescales because we want to explore the dynamic interplay of both. We therefore explicitly simulate multiple, potentially coexisting types.

We denote the different types of species *X* by *X_i_* (*i* = 1, 2, …), each characterized by its trait values *c_X_i* and *d_X_i*. The total density of species *X* at time *t* is then given by the sum of the densities of its types, defined via *x*^(*t*):= *x_i_*(*t*). Analogous considerations hold for species *Y*.

After each mutation event, the time until the next mutation event is sampled from an exponential distribution with parameter µ*_X_ · x*^(*t*), where µ*_X_* is the per-capita mutation rate for species *X* and *x*^(*t*) its total population size. At each mutation event, a “parent” type is selected with probability proportional to its relative density within the population, from which an offspring “mutant” type arises. The mutant type’s trait values are sampled from normal distributions centered on the parent type’s traits *c_X_i* and *d_X_i*, with standard deviations σ*_c_X* and σ*_d_X*, respectively. Larger values of σ*_c_X* and σ*_d_X* therefore imply greater trait variation and potential for faster evolution. We refer to these parameters as *mutation spread*. In each simulation and for each species, a value for the mutation spread σ*_c_* is fixed while the corresponding value for the mutation spread σ*_d_* is computed from the trade-off relation in Equation (4), so that a species’ combination of mutation spreads, characterizing its evolutionary strategy, is constant over time. We assume that the traits *c_X_i* and *d_X_i* are constrained by a genetic trade-off (Blows & Hoffmann 2005; Hine et al. 2014), so that fast evolution of one trait implies slow evolution of the other. Introducing a maximum mutation spread σ_max_, we implement this trade-off via

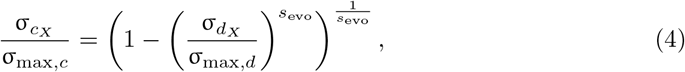

where *s*_evo_ *>* 0 determines the shape of the trade-off (Figure 1). Again, the relation is convex, concave, or linear when *s*_evo_ *<* 1, *s*_evo_ *>* 1, or *s*_evo_ = 1, respectively. After a new type’s trait values have been computed, its intrinsic growth rate *r* is immediately determined from its defence trait *d* via the ecological trade-off (Equation 3). Then, if that new type’s net growth is positive, it is introduced at density *p*, while the parent’s density is reduced by the same amount. The same procedure applies to species *Y*.

Between mutation events, all types follow ecological dynamics analogous to Equation (1), now with type-specific parameters. We hereby assume that the competitive effect of one species on the other (*b_XY_, b_Y X_*) is independent of type, to reflect the fact that these relations between intra- and interspecific competition are determined by species-specific ecological characteristics (e.g., resource-use strategies, amount of space used by each individual), which therefore don’t vary across types, even when the physiological optimum or defence traits evolve. Non-viable types are removed once their density falls below a certain threshold. We choose this threshold to be the same at which new types are introduced (*p*), so only types with a positive initial growth can enter the system (see Supporting Information S1 for details).

Finally, we implement a directional, linearly increasing environmental change *c*(*t*) at a rate τ. Consequently, if the physiological optimum of a species (*c_X_*, *c_Y_*) is fixed, the difference between the state of the environment and the physiological optimum grows linearly in time. Since the performance function approaches zero as that distance grows (see Equation (2)), the growth rate of the species will eventually be negative and its density will decline. To maintain viable densities, the species will have to evolve its physiological optimum trait.

### 2.3 Simulation experiments

To test for the effect of eco-evolutionary feedback on species survival in our study system, we design three simulation scenarios. In the *ecological scenario*, no evolution can occur. As described above, this will necessarily lead to the species’ extinction since the environmental conditions change over time. This scenario will serve as a reference. In the *1-trait evolution scenario*, only the physiological optimum can evolve. In the *2-trait evolution scenario*, both traits can evolve, subject to the trade-off in Equation (4). In the evolutionary scenarios, extinction may or may not occur, dependent on how fast evolution acts in relation to the environmental change.

To quantify the net effect of evolution and the corresponding eco-evolutionary feedback, we follow Weyerer et al. (2023) and calculate the difference in extinction times between the reference (*t_X_R*) and evolutionary (*t_X_*) scenarios, i.e., Δ*t_X_* = *t_X_ − t_X_R*. Time until extinction is hereby measured in the same unit across all simulations and used to compute delay as a comparative measure of transient dynamics. Higher absolute values of Δ*t_X_*indicate stronger eco-evolutionary feedbacks. Positive values indicate that evolution delays extinction, while negative values indicate an acceleration. When species are not extinct by the end of the simulation, we calculate this difference as Δ*t_X_* = *t_f_ − t_X_R*, where *t_f_* is the maximum simulation runtime.

In the following, we first analyze the case of a single species and then use these results to understand the more complex dynamics of two competing species. For the two-species system, we first consider the cases of single-trait and two-trait evolution with the negative ecological and evolutionary trade-offs outlined above. To test how sensitive our results are to the trade-off assumptions, we also ran simulations with positive ecological and evolutionary trade-offs, faster or slower environmental decline, as well as higher or lower mutation frequency. We explain our choice of parameters and initial conditions in the Supporting Information S2.

## 3 Results

### 3.1 Single-species system: Two-trait evolution modulates the potential for evolutionary rescue

In the absence of evolution, species eventually go extinct with increasing environmental change, but this extinction can be delayed when species are allowed to evolve (Figure 2). Besides a limited region in the convex ecological trade-off (detailed in Supporting Information S3), the curves showing extinction delay as a function of mutation spread of the physiological optimum trait are positive in both the 1-trait (dark, solid lines in Figure 2) and 2-trait (light, dashed lines in Figure 2) evolution scenarios, and across all combinations of ecological and evolutionary trade-off shapes (convex, linear, or concave). This positive effect of evolution indicates a potential for evolutionary rescue: if the change in the abiotic environment stopped somewhere between the extinction time of the reference (*t_X_R*) and the evolution (*t_X_*) scenarios, we would observe extinction in the former but survival in the latter case.

**Figure 2:**
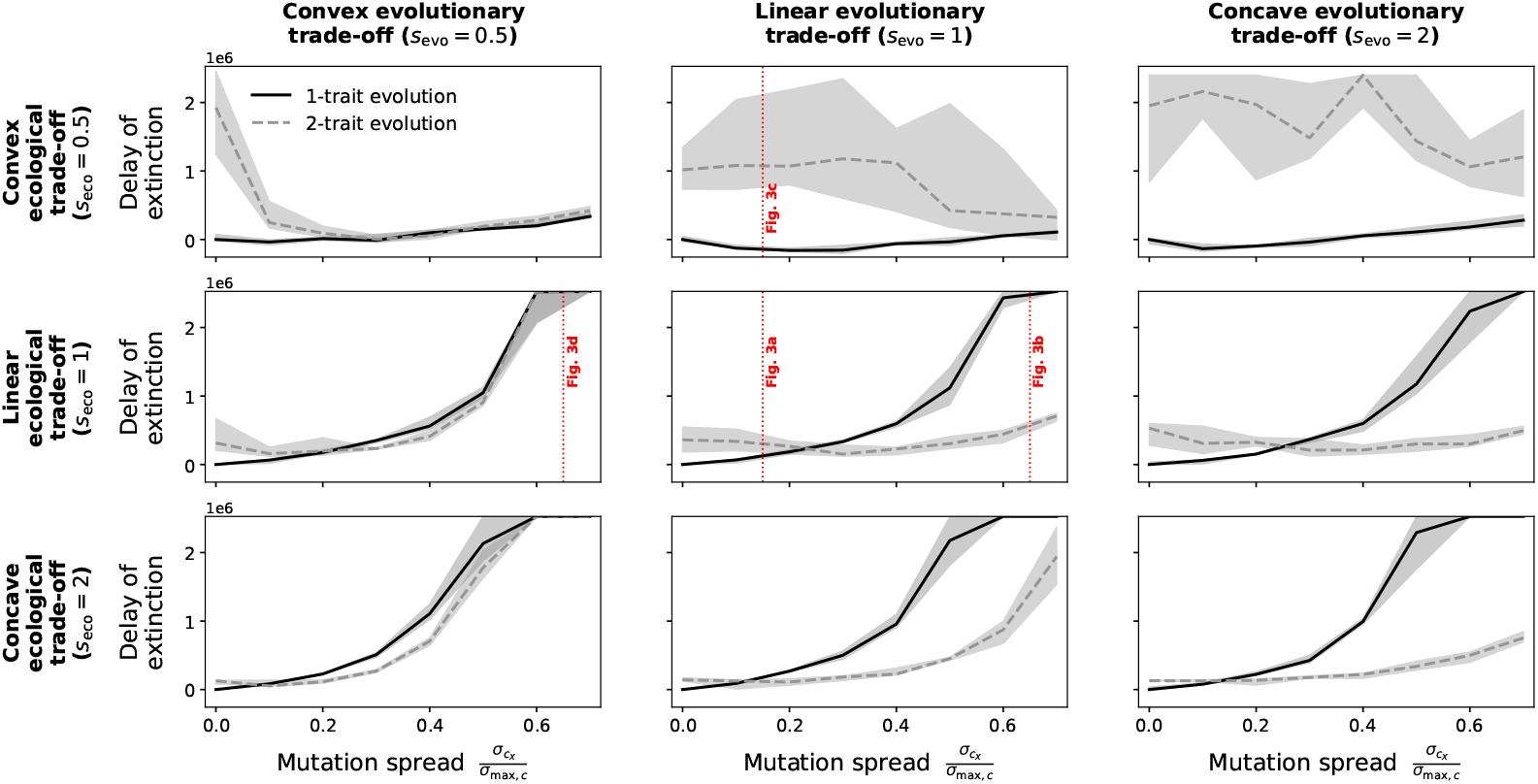
Net effect of evolution and corresponding eco-evolutionary feedback on the extinction time of a single population. The curves represent the median of the delay of extinction (Δ*t_X_* = *t_X_ − t_X_R*) as a function of mutation spread σ*_c_X* over 10 simulation runs; the shaded areas represent the interquartile range. The dark solid curves represent the extinction delay for the 1-trait evolution scenario. The lighter dashed curves represent the extinction delay for the 2-trait evolution scenario (in which evolution of *d* also implies evolution of *r* via Equation 3). Red vertical lines indicate simulation scenarios displayed in Figure 3. Unless otherwise noted, parameter values are as in Table 1.

If we focus on the linear ecological and evolutionary trade-offs (Figure 2e), we find that increasing values of the mutation spread σ*_c_X* increase the delay of the extinction time Δ*t_X_* in the 1-trait evolution scenario. This outcome was expected from eco rescue theory (REFs) and can be explained by the fact that higher values of σ*_c_X* translate into faster adaptation to the abiotic environment and hence longer survival times. Interestingly, when switching to the 2-trait evolutionary scenario, this monotonic relationship no longer holds. We find that the additional effect of evolving defence against competition on extinction delay then depends on the mutation spread: if σ*_c_X* is below some threshold, 2-trait evolution is more beneficial than 1-trait evolution; whereas if σ*_c_X* is above this threshold, 1-trait evolution is more beneficial than 2-trait evolution (solid curve above dashed curve in Figure 2e).

To explain this non-trivial outcome, we compare selected simulation runs with relatively low versus high mutation spread σ*_c_X* (Figure 3a,b). In the 2-trait evolution scenario, we generally find that defence against competition *d_X_* evolved to approach its maximum *d*_max_ = 1 (dashed red lines in Figure 3). This process can be explained through the invasion criteria (detailed in Supporting Information S4). Whether or not the evolution of defence is beneficial depends on the value of σ*_c_X*. If σ*_c_X* is small (Figure 3a), the rapid evolution to maximum defence diminishes the negative effects of competition. This results in an initial higher population density, which then buffers the population against extinction and effectively delays extinction time. However, because of the ecological trade-off, the growth rate (*r_X_*) approaches zero (dashed blue line in Figure 3a) when the defence trait approaches its maximum. If σ*_c_X* is large, it is exactly this trade-off that reduces the species’ ability to track environmental change when defence evolves (Figure 3b). First, compared to the case of small σ*_c_X*, evolution of defence against competition is slower due to the trade-off between σ*_c_X* and σ*_d_X*. Second, since evolution of high defence leads to low growth rates, new types that are better adapted to the environment will not be able to grow fast enough to ensure their persistence. This ultimately results in a too slow evolution of physiological performance, reflected by the dashed line in the third panel in Figure 3b flattening off. Hence, evolution of physiological optimum is beneficial and even required for persistence in the long run, whereas evolution of defence against competition has the short-term benefit of increasing population density and the long-term negative cost of inhibiting evolutionary change through the reduction in growth rate.

**Figure 3:**
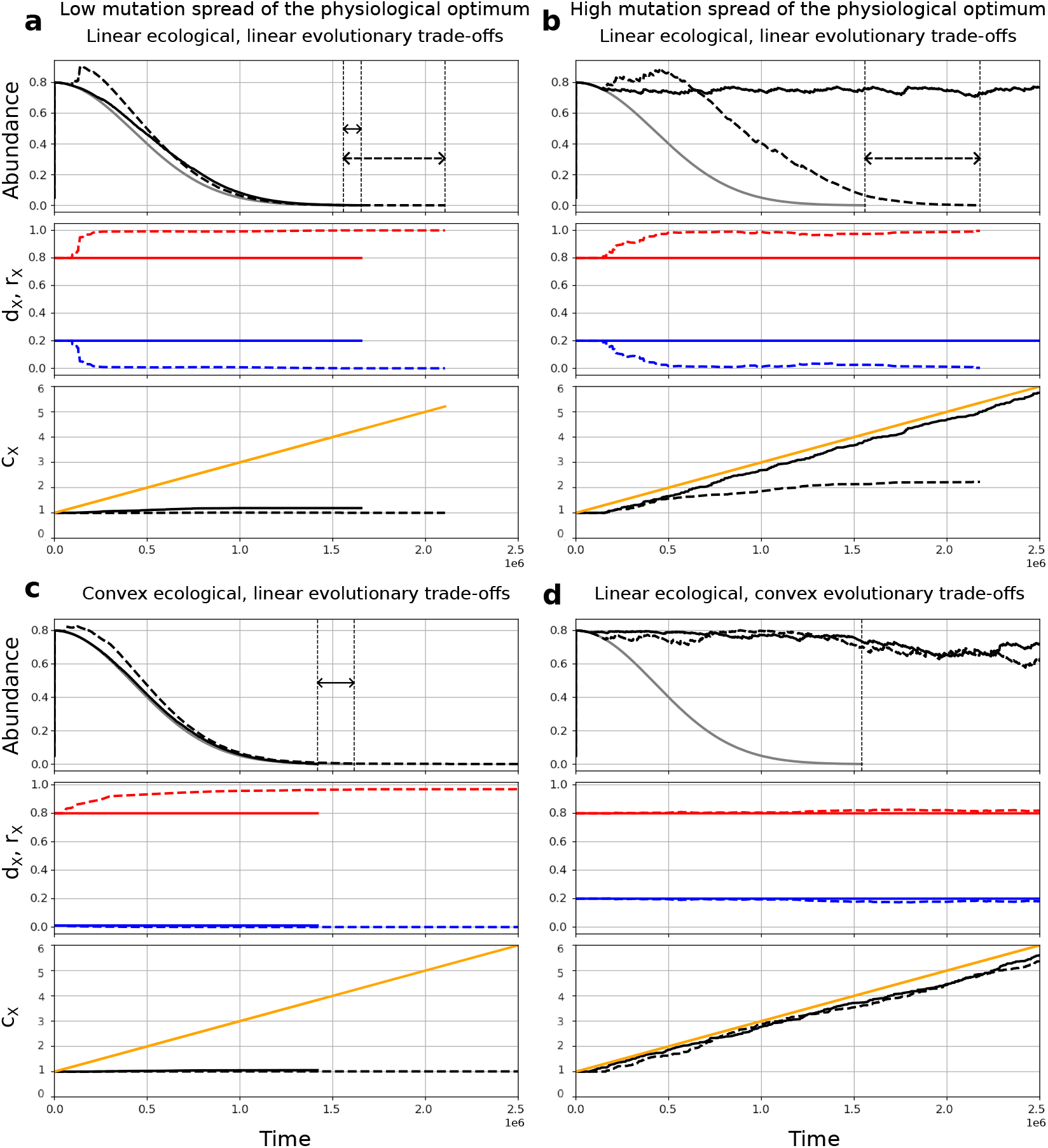
Selected simulations under low (σ*_c_X /*σ_max*,c*_) = 0.15 (a, c) and high (σ*_c_X /*σ_max*,c*_) = 0.65 (b, d) mutation spread of the physiological optimum trait, with linear-linear (a, b), convex-linear (c), and linear-convex (d) ecological-evolutionary trade-off shapes. In each of these four subfigures, panels show population density (top panel), defence against competition (red line) and growth rate (blue line) (middle panel), and environmental state (yellow) and physiological optimum (black line) (bottom panel). Solid (dashed) lines: 1-trait (2-trait) evolution; gray colors: ecological reference. Solid (dashed) horizontal arrows: extinction time delays under 1-trait (2-trait) evolution relative to the reference.

The qualitative patterns described above remain robust across ecological and evolutionary trade-off shapes. These trade-offs primarily shift the mutation spreads at which the relative benefit of 1-trait versus 2-trait evolution changes, and affect the variability across simulation runs (Figure 2). For example, changing the shape of the ecological trade-off modifies how intrinsic growth rate varies as the defence against competition evolves. This, in turn, influences population persistence by modifying the rate at which better adapted types can invade the system and the rate at which poorly adapted types decline under increasing abiotic pressure (Figure 3c). Similarly, the evolutionary trade-off determines how fast defence can evolve for a given mutation spread of the physiological optimum, which, combined with the ecological trade-off, again affects the aforementioned mechanisms (Figure 3d). In some cases, particularly under convex ecological trade-offs, stochasticity in mutation time points and trait values leads to increased variability in extinction times (see Supporting Information S5). Nevertheless, these effects do not alter the overall conclusion: in all tested scenarios of the one-species system, evolution consistently delays extinction, demonstrating a robust potential for evolutionary rescue.

### 3.2 Two-species system: competitive interactions can lead to evolutionary murder

In contrast to the single-species case, we find that evolution can delay or accelerate a species’ extinction in the two-species system, and this occurs both in the 1-trait and 2-trait evolution scenario (Figure 4a,b). Focusing on a specific case where both species only evolve one trait and differ in their mutation spread (red squares in Figure 4a, Figure 5), we observe that the faster evolving species (here *X*) is able to track the abiotic environment due to evolution, allowing it to persist and maintain high densities, while the slower evolving competitor With evolution, species *X* persists until the end of the simulation (evolutionary rescue), whereas species *Y* goes extinct much earlier, around *t ≈* 3.5 *·* 10^5^ (evolutionary murder). species (here *Y*) suffers from the increased biotic pressure in addition to increasing abiotic pressure, leading to its accelerated extinction. Importantly, this does not only reflect a relative disadvantage compared to the faster evolving species, but also a net negative effect of evolution: species *Y* goes extinct earlier than in the ecological reference scenario where neither species evolves (red squared cells in Figure 4a). This accelerated decline indicates a potential for evolutionary murder: if environmental change stopped somewhere between the extinction time point of species *Y* and the ecological reference time point *t_X_R*, we would observe persistence of both species in the ecological scenario, but extinction of species *Y* in the evolutionary scenario, due to the combined effect of environmental change and evolution of species *X*.

**Figure 4:**
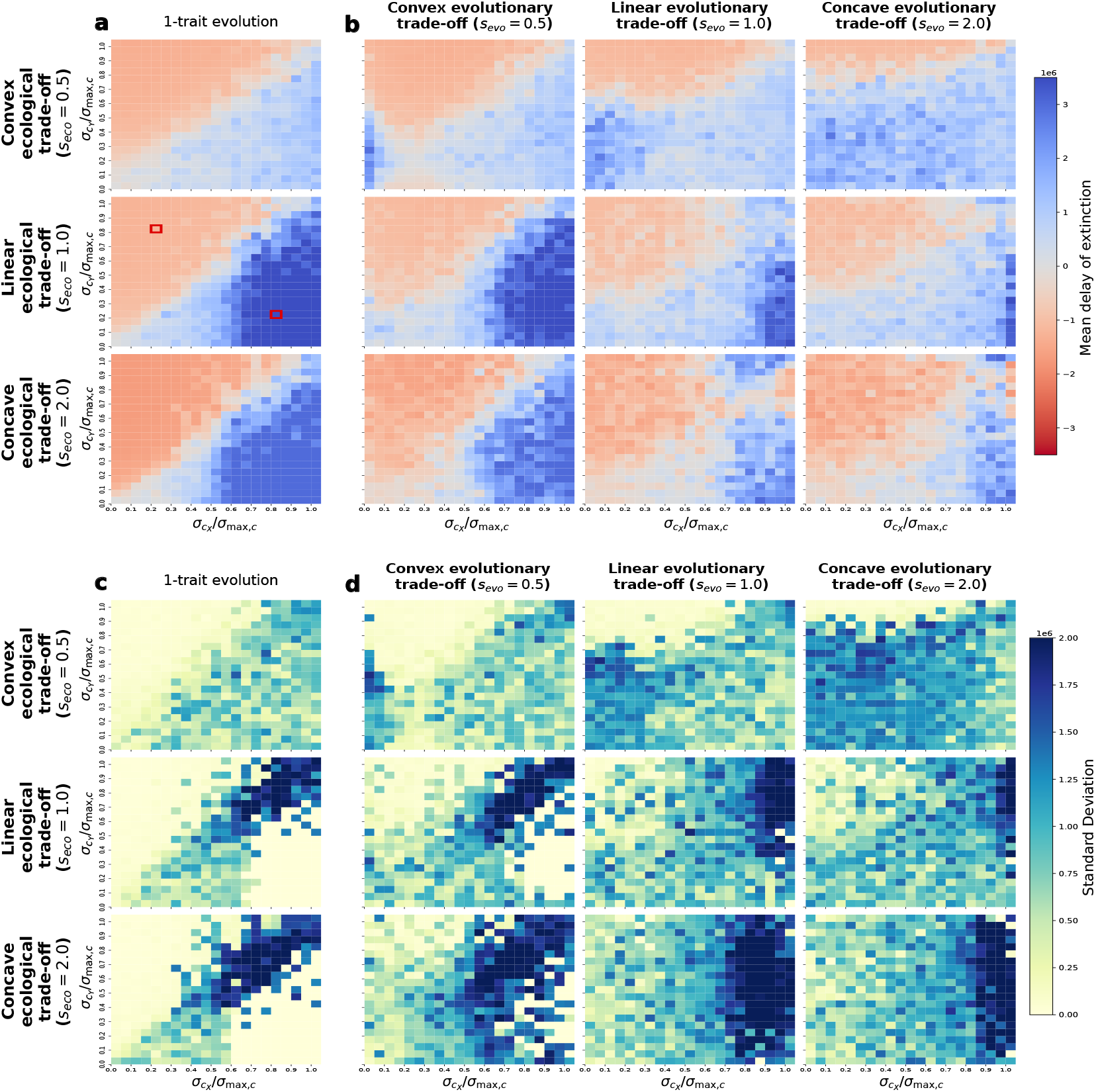
Extinction delay Δ*t_X_* (i.e., difference in extinction time between evolutionary and ecological scenarios) for species *X* in the 1-trait (a) and 2-trait (b) evolution scenario, and variability thereof over 10 replicates (c, d). In (a) and (c), since one trait evolves, no evolutionary trade-off exists, so only one heatmap is shown for each ecological trade-off shape. Colors represent (a, b) mean or (c, d) standard deviation of Δ*t_X_* across simulation runs. Positive Δ*t_X_* values (blue in a, b) indicate that evolution delays extinction of species *X*, whereas negative values (red in a, b) indicate that evolution accelerates extinction. Since species differ only in their mutation spreads, each point (*σ_c_X, σ_c_Y*) has a symmetric counterpart (*σ_c_Y, σ_c_X*): the latter corresponds to the outcome for species *Y* in the original scenario. Thus, each heatmap encodes outcomes for both species. Red squares mark cells corresponding to the simulation shown in Figure 5.

**Figure 5:**
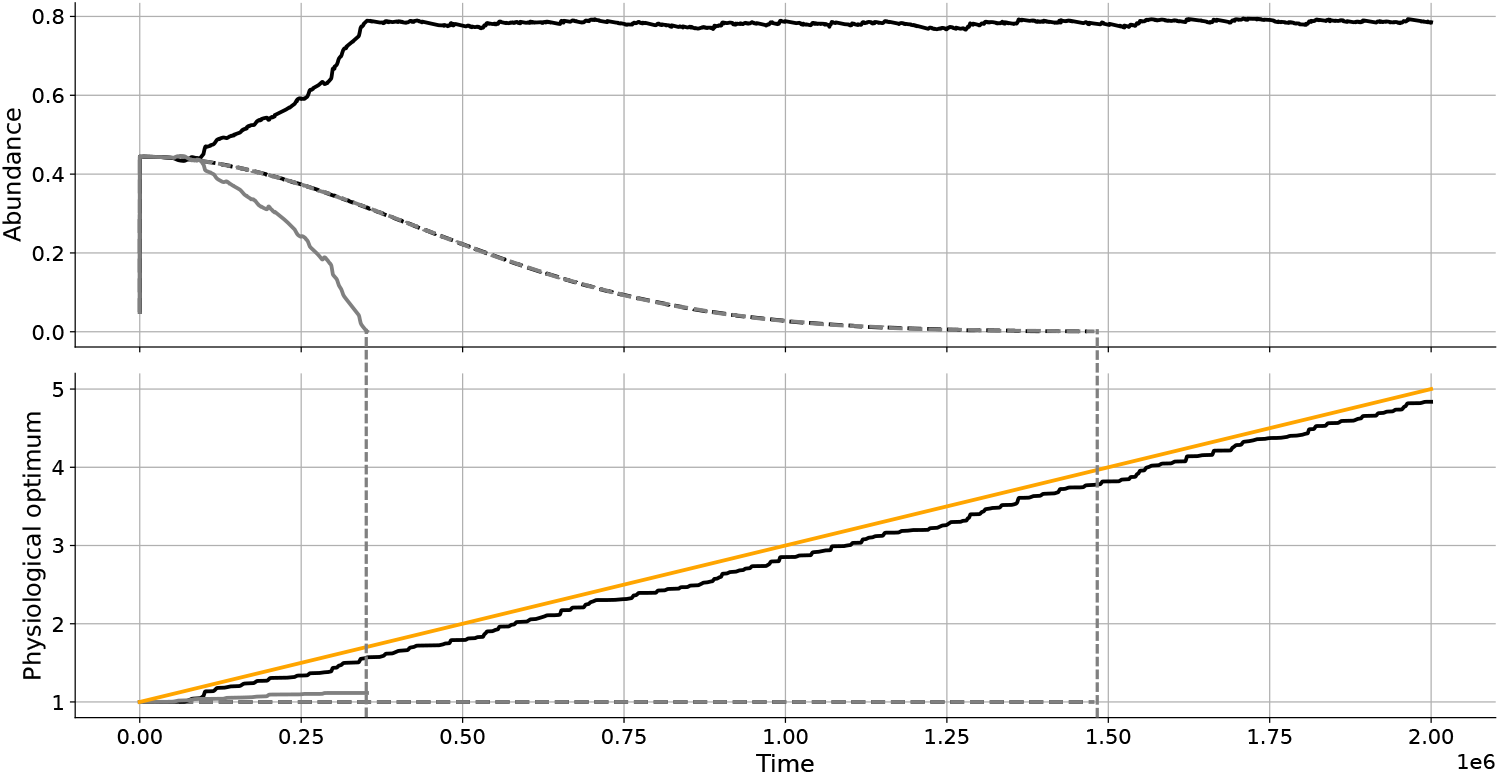
Selected simulation of two-species system, where both species evolve to the abiotic environment only and follow a negative linear ecological trade-off. (a) Species abundances of species *X* (black) and species *Y* (grey) in the reference (dashed curves) and evolution (solid curves) scenario with 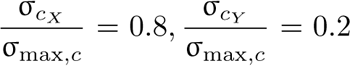. (b) Evolution of the physiological optimum for species *X* (black) and species *Y* (grey) along with the environmental state (yellow). Dashed curves correspond to the reference scenario without evolution, while solid curves represent the 1-trait evolution scenario. Vertical dashed lines indicate the timepoint of extinction. In the reference scenario, both species share the same parameters and initial conditions; therefore their dynamics are the same, and both go extinct around *t ≈* 1.5 *×* 10^6^.

This pattern of combined rescue and murder remained consistent for all ecological and evolutionary trade-off shapes (convex, linear, or concave, see Figure 4a,b), as long as the difference in mutation spread of the two species was sufficiently large. Notably, it also holds in the 2-trait evolution scenario, where faster adaptation to the abiotic environment comes at the cost of slower adaptation to competition via the evolutionary trade-off (Figure 4b). Deviations from this pattern occur mostly along the diagonal where both species have similar mutation spreads (a detailed explanation of the mechanisms is provided in Supporting Information S6). Furthermore, when testing different combinations of negative and positive ecological and evolutionary trade-offs, we observe that the described pattern of combined rescue and murder remains consistent (see Figures S1, S2, S3 in Supporting Information S7). The most striking difference between the possible combinations of negative and positive ecological and evolutionary trade-offs can be found for the convex ecological trade-off in combination with the concave evolutionary trade-off. When both are negative, a larger blue region is observed, indicating that for some parameter combinations of mutation spreads both species are rescued (Figure 4). Such pattern rapidly disappears with a positive evolutionary trade-off (Figure S1 in Supporting Information S7.1) and is reduced with a positive ecological trade-off (Figure S2 in Supporting Information S7.2). Finally, the pattern of combined rescue and murder also remains largely consistent when changing the relative time scales at which environmental and evolutionary dynamics act (Figures S4-S7 in Supporting Information S8). As expected, we find that effect sizes are overall smaller with slower evolutionary change or faster environmental change, and larger with faster evolutionary change and slower environmental change.

Consistent across all scenarios, we find that there are two reasons for high variability in our results (Figure 4c,d; Figures S1–S3 in Supporting Information S7 and Figures S4–S7 in Supporting Information S8). First, variability arises when both species track the environment at similar rates. In this case, outcomes are primarily determined by which species gains a fitness advantage first (reflecting stochasticity in the timing of mutation events), leading to variability concentrated along the diagonal and increasing with larger mutation spreads (Figure 4c). Second, variability is high in regions where evolution strongly delays extinction (dark blue region in Figure 4a,b; Figures S1–S3 in Supporting Information S7 and Figures S4–S7 in Supporting Information S8). This is due to the fact that extinction times can vary widely with positive delays, that are, in principle, unbounded, whereas advances in extinction are constrained by the baseline ecological scenario. As a result, variability is typically higher in regions of strong evolutionary rescue than in regions of evolutionary murder. Note that this is not always visible in the heatmaps because of the fixed simulation runtime: in cases when evolutionary rescue is strong, the focal species persists until the end of a simulation, *t_f_*, across all replicates, so extinction times do not vary. With longer simulation times, variability would likely emerge in these regions.

## 4 Discussion

Considering both abiotic and biotic changes when studying ecological and evolutionary dynamics is crucial given that both theoretical (Johansson 2008; Northfield & Ives 2013; Drury et al. 2018; Loeuille 2019; Wang et al. 2025) and empirical studies (Yoshida et al. 2003; Hart et al. 2019; Govaert et al. 2021; Grainger et al. 2021) have shown that biotic interactions can induce rapid evolution and alter evolutionary trajectories of species in response to abiotic change. Considering ecological and evolutionary dynamics simultaneously is thus important if we aim to understand and predict species responses to global change (Norberg et al. 2012; Urban et al. 2024). However, whether evolution will be beneficial or detrimental in a multi-species setting is not straightforward. Our eco-evolutionary competition model tracked the ecological densities and evolutionary trait dynamics of one or two competing species exposed to a directional abiotic environmental change. Using this model, we show that evolution is generally beneficial in the single-species system, while it holds potential for evolutionary murder in a 2-species system. For both systems, these results were largely independent of the considered ecological and evolutionary trade-off shapes (convex, linear, or concave). Even more so, for the 2-species system, patterns of evolutionary rescue and murder remained remarkably consistent among combinations of negative or positive ecological and evolutionary trade-offs, and also when considering slower or faster relative timescales of adaptation and environmental decline. In the following, we discuss our findings in more detail, starting with the single-species system.

In the single-species model, 1-trait and 2-trait evolution were overall beneficial for survival, although the extent of extinction delay depended on the number of evolving traits and on the combination of the ecological and evolutionary trade-off shapes. When only the physiological optimum trait evolved, increasing the mutation spread progressively delayed extinction. This result is in line with previous work on evolutionary rescue (Gomulkiewicz & Holt 1995; Carlson et al. 2014), demonstrating that larger genetic diversity (e.g., in microbial species *Pseudomonas fluorescens*; Ramsayer et al. 2013) and greater population size (e.g., in yeast *Saccharomyces cerevisiae*; Bell & Gonzalez 2009) increase the potential for evolutionary rescue. Within a population, multiple traits can be genetically correlated, so that the evolution of a trait can be restricted by genetic or evolutionary constraints (Hine et al. 2014). When two traits evolved in our model, the delay of extinction was not always greater than when only the physiological optimum evolved, indicating the observation that additional evolving traits do not necessarily enhance evolutionary rescue (Campbell et al. 2025). Nevertheless, both 1-trait and 2-trait evolution consistently extended persistence compared to the reference scenario.

Contrary to the single-species setting, evolution did not always lead to longer persistence in the two-species system, because rescue of one species often led to evolutionary murder of its competitor. Evolutionary murder occurred when one species could track the environmental change considerably faster than its competitor, establish high population densities and exert increased competitive pressure. This reduced the competitor’s density and hence its evolutionary potential to track the environmental change, thereby accelerating its decline. This mechanism coincides with previous observations (Jones 2008), which found that competition can prolong persistence of a faster-evolving species, hereby shortening that of its competitor. By explicitly comparing extinction timepoints in scenarios with and without evolution, our study extends this finding and systematically demonstrates evolutionary rescue and murder in 2-species competitive systems across all combinations of ecological and evolutionary scenarios considered here. Despite finding some differences in extinction patterns between ecological and evolutionary trade-offs, the overall pattern of evolutionary rescue and murder is remarkably robust. This robustness of our results indicates that evolutionary murder might be common in competitive systems and calls for further research into the interaction of ecological and evolutionary processes and their joint effect on species persistence.

To the best of our knowledge, empirical evidence of evolutionary murder remains almost absent (but see Greenrod et al. 2026), perhaps because it can resemble classical competitive exclusion, which does not involve evolutionary processes (Hardin 1960). By using ecological and evolutionary quantities that can be tracked in empirical systems, our study provides practical guidance on how evolutionary rescue and murder could be quantified in experimental settings. For competitive systems, empirical studies could quantify defence against competition by measuring changes in individual performance (e.g., comparing biomass or growth) as a function of competitor density, yielding growth-response metrics that capture the sensitivity of a species to such biotic pressure (Goldberg & Landa 1991; Semchenko et al. 2018). Common garden or reciprocal transplants would hereby be adequate experiments to attribute such changes in performance to evolution. Similarly, traits describing the response to abiotic conditions can be inferred from curves relating individual performance, such as growth, to environmental variables, where the location of maximal performance defines the optimal trait value (Childress & Letcher 2017). Other approaches might involve combining evolved with non-evolved populations of different competitor species and tracking their density dynamics over time. At a minimum, this would first require creating a setup of populations of at least two species evolved to an abiotic environment, interspecific competition and their combination. Second, these evolved populations could then be set out to compete against one another. By quantifying differences in their density dynamics with and without evolved competitor could then demonstrate indirect evolutionary effects (either rescue or murder). Alternatively, experimental practices to halt evolution, such as replacing evolving populations by ancestral individuals repeatedly throughout an experiment (Hart et al. 2019), could also reveal patterns of evolutionary murder. Possible experimental model systems include not only bacteriophages (Greenrod et al. 2026) or ciliates (Govaert et al. 2021), but also phytoplankton (Ŕeveillon & Becks 2024), zooplankton (Vanvelk et al. 2024), duckweed (Hart et al. 2019), and plant weeds (Alexander et al. 2015), which have generation times suitable for observations of the effect of eco-evolutionary interactions in experiments. Finally, our results have important implications for systems with more species. Several recent studies suggest that the concept of evolutionary rescue can be extended towards multi-species systems (e.g., Fussmann & Gonzalez 2013; Low-Décarie et al. 2015; Yamamichi & Miner 2015; Bell et al. 2019; Fugère et al. 2020; Yacine et al. 2021; van Velzen 2023). A large proportion of these studies focuses on trophic interactions, demonstrating for example that prey adaptation to external stressors can enable predator survival (Yamamichi & Miner 2015; van Velzen 2023). By contrast, our model predicts that biotic interactions within multi-species systems can not only reduce the potential for evolutionary rescue, but also enable evolutionary murder. This is in line with insights from the predator-prey food web model by van Velzen (2023), where evolutionary murder could occur within trophic levels, that is, between species that interact indirectly via resource competition (sharing the same prey) or apparent competition (sharing the same predator). Our results are also consistent with a spatially explicit eco-evolutionary model of multi-species responses to climate change predicting that competitive interactions can lead to extinction debts (Norberg et al. 2012) and with an individual-based competition model demonstrating that community-wide rescue can lead to rapid loss of rare species (van Eldijk et al. 2020). These studies and our results suggest that whether evolution is beneficial or detrimental for species persistence in multi-species ecological networks could depend on the type of biotic interaction dominating these networks, with antagonistic systems tending towards rescue and competitive systems tending towards evolutionary murder and hence partial network collapse. We argue that further empirical and theoretical research is needed to test how networks composed of multiple interaction types, as conceptually outlined by Fontaine et al. (2011), can help us understand and forecast eco-evolutionary responses in more realistic systems.

## Acknowledgements

We thank the whole Eco-Evolutionary Modelling group at the University of Hohenheim for insightful discussions and moral support. In particular, we thank Felix Jäger and Dimitrios Nakos for a code review and for helpful feedback on the manuscript. The corresponding author acknowledges invaluable support of Antoine and Luna.

## Data and code availability statement

Code for reproducing our results is available via Zenodo: https://doi.org/10.5281/zenodo.17982254

## Author contribution statement

SL: Code development, Conceptualization, Investigation, Methodology, Writing - original draft, Writing - review and editing

FL: Conceptualization, Investigation, Writing - review and editing

KTA: Conceptualization, Investigation, Methodology, Supervision, Writing - original draft, Writing - review and editing

LG: Conceptualization, Investigation, Funding acquisition, Supervision, Writing - original draft, Writing - review and editing.

