## Supporting Information for "Eco-evolutionary dynamics in competitive systems: Rescue and murder"

### Contents

**S1 Type-specific ecological dynamics** **3**

**S2 Choice of model parameters** **3**

**S3 Explanation for accelerated extinction in the 1-species system** **5**

**S4 Evolution of defence against competition** **6**

**S5 Variability in extinction delay observed in the 1-species system with 2-trait evolution under convex ecological and concave evolutionary trade-offs** **7**

**S6 Effects of different trade-off shapes on the eco-evolutionary outcomes observed in the 2-species system under negative trade-offs** **9**

**S7 Robustness of our results in the 2-species systems, relaxing the assumption of negative trade-offs** **12**

**S8 Relative timescales of ecology, evolution and environmental change** **18**

**S9 Stable coexistence for the initial types** **23**

### S1 Type-specific ecological dynamics

Between two mutation events, the existing types follow the same dynamics as given in Equation (1) of the main text, modified for multiple types as follows:

$$\begin{cases} \dfrac{\mathrm{d}x_i}{\mathrm{d}t} = r_{X,i}x_i\left(w_{X,i}(c(t)) - a_{X,i}(\hat{x} + b_{XY}\hat{y})\right), \\ \dfrac{\mathrm{d}y_j}{\mathrm{d}t} = r_{Y,j}y_j\left(w_{Y,j}(c(t)) - a_{Y,j}(\hat{y} + b_{YX}\hat{x})\right), \end{cases} \tag{S1.1}$$

where $\hat{x}(t) = \sum x_i(t)$ and $\hat{y}(t) = \sum y_i(t)$ are the total densities of species $X$ and $Y$ at time $t$, respectively. Note that the number of types changes after a mutation event and can differ between the two species, depending on how many mutation events have occurred for each species. The parameters are now type-specific as indicated by the subscripts $i$ and $j$; their meaning is as in Equation (1) in the main text; see Table 1 in main text. We assume that the ratio between sensitivity to intra- and interspecific competitive pressure $(b_{XY}, b_{YX})$ is independent of type. Solutions of the system given in Equation (S1.1) with positive initial conditions remain positive, but the density of a non-viable type will approach zero. To avoid unrealistically low densities, we set the densities to zero for all types whose density falls below $p$, the same density at which new types are introduced. With this choice, only types with positive initial growth can enter the system.

### S2 Choice of model parameters

Since we aim to observe evolution in response to environmental change, we choose the pace of abiotic change to be slow (i.e., small value of $\tau$) relative to the potential speed of evolutionary adaptation. To ensure the initial survival of populations, we set the initial physiological optima to match the initial state of the environment ($c_X = c_Y = c(0)$). Since mutations are rare, we choose the per capita mutation rates ($\mu_X, \mu_Y$) and the density at which new mutant types are initialized ($p$, which is also the extinction threshold) to be very small compared to unit time and to the initial and maximum population density.

In simulations where both species are initially present, we choose parameters such that the two competitors can stably coexist at the initial state of the environment, whereby we assume that the spread of the physiological performance function is the same for both species (see Supporting Information S9 for details). We also choose the initial values of $d_X, d_Y, b_{XY}$ and $b_{YX}$ to be equal (see Table 1 in the main text) such that the two species are initially identical and only differ in their mutation spreads. That way, all differences in outcome are due to evolution.

#### S3 Explanation for accelerated extinction in the 1-species system

In Figure 2 in the main text, the delay in extinction between the reference and evolution scenarios is shown as a function of the mutation spread of physiological performance, $\sigma_{c_X}$. We generally expect this delay to increase with $\sigma_{c_X}$, as a larger mutation spread allows for larger changes in the trait values of the physiological optimum and hence could result in faster evolution. This faster evolution should enable the species to better track environmental change and thus persist longer. However, at low values of $\sigma_{c_X}$, our model shows earlier extinction in the evolution case than in the reference one. This counterintuitive result does not reflect a biological process, but rather a numerical consequence of our model implementation.

In our simulations, evolution proceeds through the introduction of successive new types generated from existing types of a species. When evolution is allowed, multiple types can therefore coexist at any given time. To ensure numerical feasibility, we imposed an extinction threshold $p$ (that is also the density of a new type in the population), such that when the density for any type falls below $p$, it is removed from the system. This threshold is applied at the type level, not at the species level, thus preventing very high numbers of coexisting types in the system which could make simulations computationally intractable. As the total population declines toward extinction, several types may independently fall below $p$ and go extinct. This sequential loss of types causes abrupt reductions in total abundance, accelerating extinction relative to the reference case, in which the entire population consists of a single (the original) type and is thus less likely to fall below the threshold. The resulting earlier extinction in the evolving scenario is thus a consequence of implementing the extinction threshold at the level of types.

#### S4 Evolution of defence against competition

In our model, a new type can invade the system if its initial growth rate is positive. From Equation (5) in the main text, we see that, for type $i$ of species $X$ at time $t$, this condition translates into

$$w_{X,i}(c(t)) \cdot d_{X,i} > \hat{x}(t) + b_{XY}\hat{y}(t), \tag{S4.1}$$

where $\hat{x}$ and $\hat{y}$ are the total population densities of species $X$ and $Y$, respectively.

Note that, in general, the invasion condition of a type (Equation (S4.1)) depends on both of its trait values: the physiological optimum trait $c_X$ (through the performance function $w_X$) and the defence trait $d_X$. However, if only $d_X$ is allowed to evolve, then evolution will favor higher values of $d_X$.

Evolution of higher defence is also predicted by the theory of adaptive dynamics (Metz et al. 1995) in a very simple logistic growth model. The resident type satisfies the equation

$$\frac{\mathrm{d}x_R}{\mathrm{d}t} = r_R x_R^* \left(w_R - a_R x_R^*\right),$$

which gives an equilibrium value of $x_R^* = w_R d_R$. A mutant type can then invade if its net growth rate at the resident-only equilibrium, given by

$$\frac{\mathrm{d}x_M}{\mathrm{d}t} = r_M x_M \left(w_M - a_M x_R^*\right) \tag{S4.2}$$

is positive. This quantity (right-hand side in Equation (S4.2)) is positive if

$$w_M d_M > w_R d_R.$$

Hence, if defence is the only trait that evolves, then the mutant type can invade if and only if $d_M > d_R$. In our study, defence against competition does not evolve in isolation, but the condition in Equation (S4.1) nevertheless clarifies why we generally observe increasing defence, consistent with predictions from adaptive dynamics theory.

### S5 Variability in extinction delay observed in the 1-species system with 2-trait evolution under convex ecological and concave evolutionary trade-offs

The variability in extinction times across simulations arises from subtle differences in the rate at which the population of the last surviving type declines. All else being equal, a type with a lower intrinsic growth rate $r_{X,i}$ will decline slower when exposed to increasing abiotic pressure, since $r_{X,i}$ scales the rate of decline in Equation (S1.1). It will therefore persist longer before extinction than one with a higher $r_{X,i}$. This effect makes the extinction time highly dependent on the final value of $r_{X,i}$, which, in turn, is shaped by the evolutionary trajectories of the defence against competition up to that point. In particular, the curvature of the evolutionary trade-off function influences both the speed and the variability of evolution to the biotic pressure. Under a concave evolutionary trade-off, this rapid evolution occurs when the mutation spread $\sigma_{c_X}$ is low, and thus $\sigma_{d_X}$ is high. However, the resulting large mutational changes can have a detrimental effect: as the mean trait value $d$ approaches its upper limit (i.e., $d \approx 1$), new types are increasingly unlikely to enter the population because many sampled values either exceed the viable range (meaning $d > 1$ and therefore $r < 0$) or the invasibility criterion (Equation (S4.1)) is not met because the value of $d$ is too small.

Consequently, the population can become “trapped” at a suboptimal final $d$ value. In such cases of concave evolutionary trade-off with suboptimal final value of $d$, the corresponding relatively high value of $r$ leads to faster decline and earlier extinction. Conversely, when $d$ evolves closer to its maximum, the final value of $r$ is lower and the population decays more slowly, resulting in later extinction. The dependence of the final values of $d$ and $r$ on the previously introduced types’ trait values and the stochasticity of the trait value sample procedure in cases of convex-concave ecological-evolutionary trade-off thus explains the highly variable extinction time point between simulation runs observed in Figure 2 of the main text.

#### S6 Effects of different trade-off shapes on the eco-evolutionary outcomes observed in the 2-species system under negative trade-offs

In the 1-trait evolution scenario, independent of the ecological trade-off, species $X$ benefits from evolution when its mutation spread is larger than that of its competitor ($\sigma_{c_X} > \sigma_{c_Y}$, blue region in Figure 4a in the main manuscript) and suffers when it is smaller ($\sigma_{c_X} < \sigma_{c_Y}$, red region Figure 4a). When $\sigma_{c_X}$ is larger than $\sigma_{c_Y}$, species $X$ adapts faster to the changing environment, maintaining a higher abundance. Therefore, species $Y$ experiences increased abiotic stress and is exposed to biotic pressure for longer, leading to its earlier extinction. The same reasoning holds when $\sigma_{c_X}$ is lower than $\sigma_{c_Y}$, but now results in $X$ going extinct earlier than in the ecological scenario. When $\sigma_{c_X} \approx \sigma_{c_Y}$, the difference in extinction times is close to zero (pale colors along the diagonal, Figure 4a), but the variability in outcomes is high (Figure 4c). With comparable mutation spreads, both species track the environmental change at comparable speeds. Hence, the stochastic timing of mutations and mutant trait values determine which species first gains a significant advantage. For the convex ecological trade-off, the net effect of evolution is less pronounced (i.e., paler red and blue regions) due to lower initial growth rates (Figure 1 in the main manuscript), which reduce the beneficial effect of evolution.

In the 2-trait evolution scenarios with linear and concave ecological trade-offs (panels in the middle and bottom row of Figure 4b in the main text), the colors are paler than in the 1-trait evolution scenario, indicating long-term costs of rapid evolution to larger values of defence against competition. Although increased mutation spread of the physiological optimum trait improves environmental tracking, this effect is counteracted by evolution to biotic interactions, which favors higher defence $d$ at the expense of a lower intrinsic growth rate $r$. The resulting reduction in $r$ slows the invasion of fitter types and limits adaptive responses, so species generally struggle to keep pace with environmental change, leading to shorter persistence times than in the 1-trait evolution scenario, although persistence until simulation end occurs in a few replicates under convex evolutionary trade-off. These compensatory effects smooth the transition between cases where one or the other species persists longer, replacing the sharp red–blue boundary of the 1-trait evolution scenario with a more diffuse gradient and inducing more variability when species differ strongly in mutation spread of defence (enlarging of the green-blue area observed outside the diagonal in Figure 4d in the main text). However, under linear and concave evolutionary trade-off and for high values of $\sigma_c$, both species can still track environmental change effectively despite reduced $r$. In this regime, outcomes become governed by biotic interactions: the species with higher defence excludes its competitor, producing the reversed red–blue pattern observed in the upper-right regions of the corresponding heatmaps (middle and bottom rows, middle and right columns in Figure 4b in the main text).

By contrast, a convex ecological trade-off (panels in top row of Figure 4b in the main text) reduces the potential for evolutionary murder, strongly diverging from the 1-trait evolution scenario. This reduction is more prominent as the evolutionary trade-off changes from convex to linear and from linear to concave. Under a convex ecological trade-off, initial growth rates are low (Figure 1 main text), and the invasion of better-adapted mutant types is slow, so both species fail to track the shifting environment. The dominant evolutionary change is therefore the increase in defence against competition. With a linear evolutionary trade-off, this evolution proceeds rapidly for species $X$ as $\sigma_{c_X}$ decreases, while under a concave trade-off it occurs rapidly across nearly all $\sigma_{c_X}$ values. Higher $d$ values temporarily release species $X$ from competitive pressure, allowing an early population increase that delays extinction. Although both populations soon decline as abiotic stress intensifies, the transient competitive release produces overall longer persistence. Even when $\sigma_{d_Y}$ is large, the additional competition arising from species $Y$'s increase in abundance remains negligible, since both species quickly lose fitness to the changing environment. This mechanism explains the predominantly blue pattern in the upper-right panel of Figure 4b. Note that, for the same reason as in the 1-species system, the combination of convex ecological and concave evolutionary trade-off increases variability in the timing of extinction of species $X$ (detailed Supporting Information S5), explaining the dark green shades observed in the top right panel of Figure 4d.

### S7 Robustness of our results in the 2-species systems, relaxing the assumption of negative trade-offs

Here, we demonstrate that our main findings of evolutionary rescue and evolutionary murder remain robust along different combinations of positive and negative ecological and evolutionary trade-offs.

#### S7.1 Effect of a positive evolutionary trade-off

In the main text, we assume a negative evolutionary trade-off between $\sigma_c$ and $\sigma_d$, such that increasing $\sigma_c$ reduces $\sigma_d$ (Figure 1, main text). Here, we consider a *positive* evolutionary trade-off, where higher $\sigma_c$ values correspond to higher $\sigma_d$, following the formula:

$$\sigma_d = \sigma_{\max,d}\left[1-\left(1-\left(\frac{\sigma_c}{\sigma_{\max,c}}\right)^{s_{\text{evo}}}\right)^{\frac{1}{s_{\text{evo}}}}\right], \tag{S7.1}$$

and a negative ecological trade-off. The positive evolutionary trade-off implies that faster evolution in response to abiotic pressure (via $c$) is now coupled with faster evolution in response to biotic pressure (via $d$). We find that the main effect of considering such a positive evolutionary trade-off is that the results from the 2-trait evolution scenario now resemble the results from the 1-trait evolution scenario. More precisely, the observed pattern of combined rescue and murder now occurs not only for species that differ substantially in their mutation spreads but also for species with more similar mutation spreads, i.e., close to the diagonal on panel b of Figure S1, due to the strong advantage a species has when evolving both traits faster than its competitor.

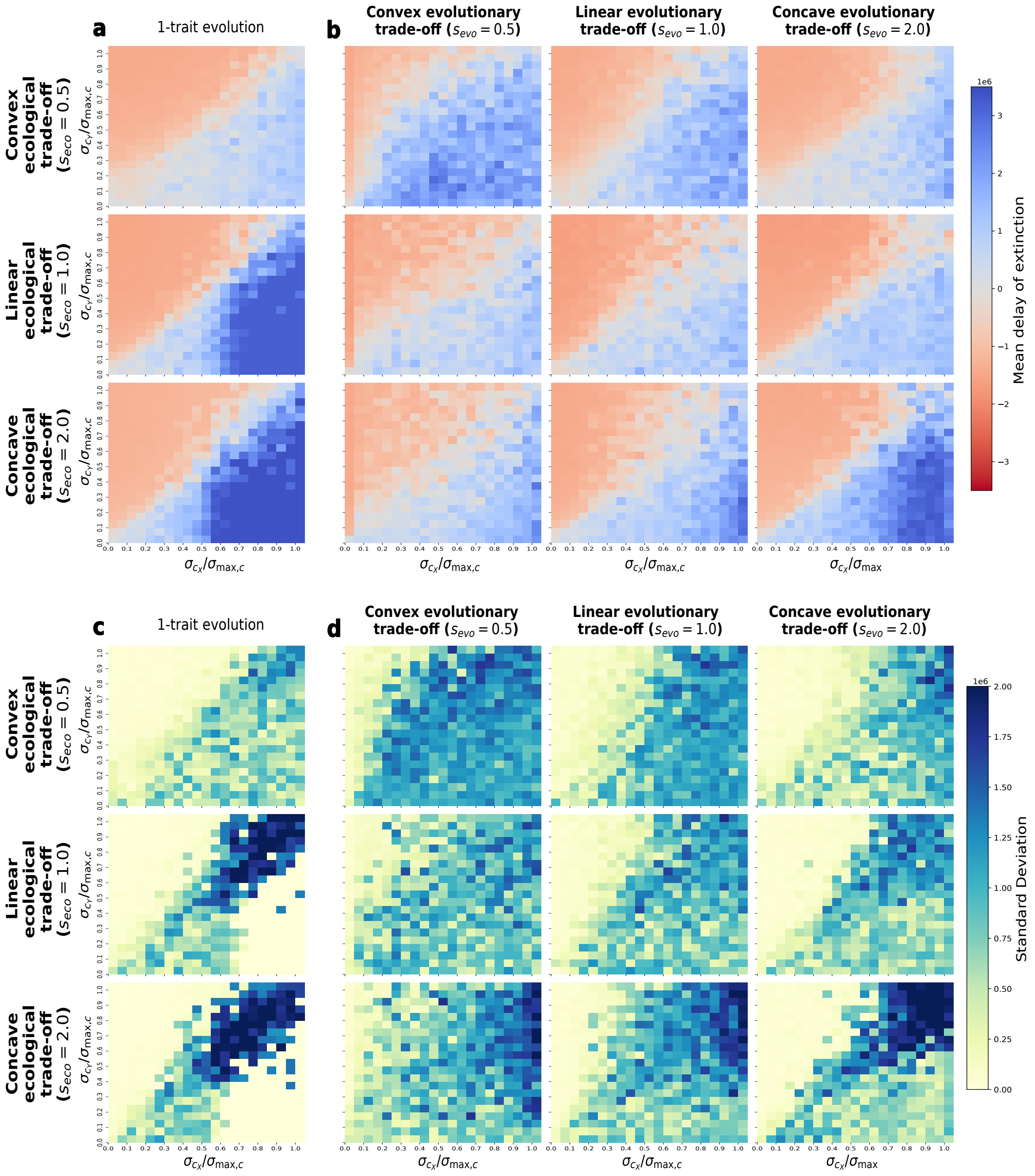

Figure S1: Extinction delay $\Delta t_X$ (i.e., difference in extinction time between evolutionary and ecological scenarios) for species $X$ in the 1-trait (a) and 2-trait (b) evolution scenario, and variability thereof over 10 replicates (c, d), under positive covariance of $\sigma_d$ and $\sigma_c$ and negative covariance of $d$ and $r$. In (a) and (c), given that only one trait evolves, no evolutionary trade-off exists, so only one heatmap is shown for each ecological trade-off shape. Colors represent (a, b) mean or (c, d) standard deviation of $\Delta t_X$ across 10 replicated simulation runs. Positive $\Delta t_X$ values (blue colors in a, b) indicate that evolution delays the extinction of species $X$, whereas negative values (red colors in a, b) indicate that evolution accelerates extinction.

#### S7.2 Effect of a positive ecological trade-off

In the main text, we assume a negative ecological trade-off between $r$ and $d$, such that increasing $d$ reduces $r$ (Figure 1, main text). Here, we consider a *positive* ecological trade-off, where $r$ increases with increasing $d$, following the formula:

$$r = r_{\max}\left[1 - \left(1 - \left(\frac{d}{d_{\max}}\right)^{s_{\mathrm{eco}}}\right)^{\frac{1}{s_{\mathrm{eco}}}}\right], \tag{S7.2}$$

and a negative evolutionary trade-off, meaning that larger values of $\sigma_c$ result in lower values of $\sigma_d$. A notable difference to the results using negative ecological trade-off arises when both species have large values of $\sigma_c$ so that they both can track the environmental change. Here, the species with a higher value of $\sigma_c$, and thus a lower value of $\sigma_d$, increases its defense slower and, due to the positive ecological trade-off, also increases its intrinsic growth rate slower than its competitor. It is this difference in lower defence and growth rate that results in a species getting outcompeted (red region in Figure S2). Overall, the qualitative outcome of extinction delays remains primarily affected by the species' ability to respond to the increasing abiotic pressure. Regions of variability in extinction delay also increase compared to the negative ecological trade-off scenario. In particular, for high $\sigma_{c_X}$ values, we observe larger (potential for) variability with a decreasing competitor's value $\sigma_{c_Y}$ (Figure S2d). In such a scenario, although the competitor evolves through smaller mutational steps to the abiotic environment, the associated increase in intrinsic growth rate under the positive ecological trade-off partly compensates for this reduction, allowing it to effectively track the environment and maintain substantial persistence. Based on the stochasticity of mutation events, there is increased variability in the subsequent increase of biotic pressure on the focal species and its extinction delay.

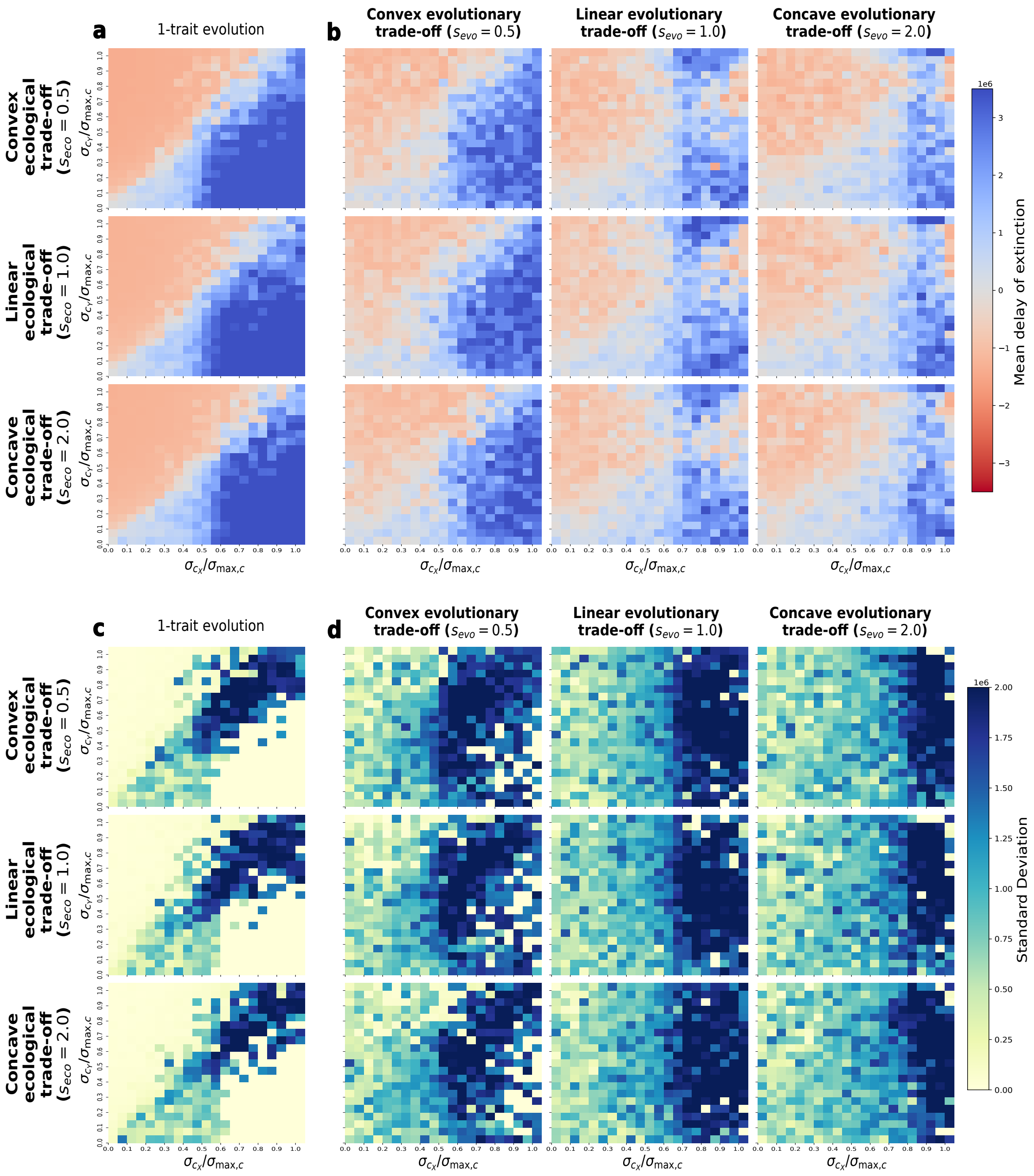

Figure S2: Extinction delay $\Delta t_X$ (i.e., difference in extinction time between evolutionary and ecological scenarios) for species $X$ in the 1-trait (a) and 2-trait (b) evolution scenario, and variability thereof over 10 replicates (c, d), under positive covariance of $d$ and $r$ and negative covariance of $\sigma_d$ and $\sigma_c$. In (a) and (c), given that only one trait evolves, no evolutionary trade-off exists, so only one heatmap is shown for each ecological trade-off shape. Colors represent (a, b) mean or (c, d) standard deviation of $\Delta t_X$ across 10 replicated simulation runs. Positive $\Delta t_X$ values (blue colors in a, b) indicate that evolution delays the extinction of species $X$, whereas negative values (red colors in a, b) indicate that evolution accelerates extinction.

##### S7.3 Effect of combined positive evolutionary and ecological trade-offs

When we consider both a positive evolutionary and a positive ecological trade-off, we obtain results similar to the ones found of a positive evolutionary and negative ecological trade-off, indicating that the effect of different evolutionary trade-offs outweighs the effect of different ecological trade-offs. Hence, the patterns observed in Figure S3 follow the same mechanisms as described in subsection S7.1.

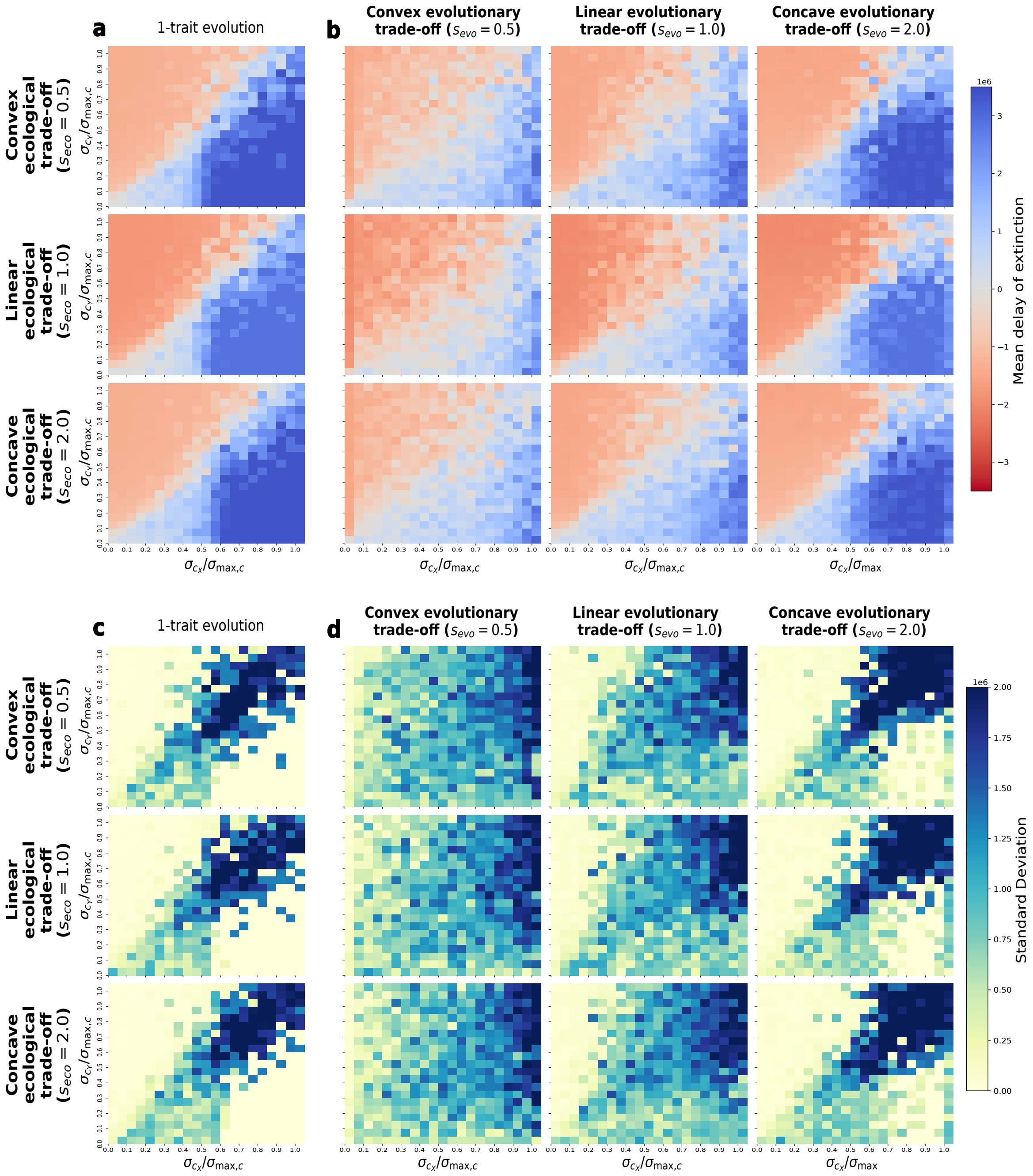

Figure S3: Extinction delay $\Delta t_X$ (i.e., difference in extinction time between evolutionary and ecological scenarios) for species $X$ in the 1-trait (a) and 2-trait (b) evolution scenario, and variability thereof over 10 replicates (c, d), under positive covariance of $d$ and $r$ as well as between $\sigma_d$ and $\sigma_c$. In (a) and (c), given that only one trait evolves, no evolutionary trade-off exists, so only one heatmap is shown for each ecological trade-off shape. Colors represent (a, b) mean or (c, d) standard deviation of $\Delta t_X$ across 10 replicated simulation runs. Positive $\Delta t_X$ values (blue colors in a, b) indicate that evolution delays the extinction of species $X$, whereas negative values (red colors in a, b) indicate that evolution accelerates extinction.

### S8 Relative timescales of ecology, evolution and environmental change

To assess whether changes in evolutionary and environmental timescales affect our results, we ran additional simulations in which we varied the speed at which evolutionary change (by modifying the mutation rate $\mu$) and environmental change (by modifying $\tau_c$ could occur. More specifically, we ran simulations with both slower ($\mu_X = \mu_Y = 5 \cdot 10^{-5}$, Figure S4) and faster ($\mu_X = \mu_Y = 1.5 \cdot 10^{-3}$, Figure S5) mutation rates compared to the main scenarios ($\mu_X = \mu_Y = 5 \cdot 10^{-4}$) presented in the main text. Similarly, we investigated how a slower ($\tau_c = 1 \cdot 10^{-6}$, Figure S6) or faster rate ($\tau_c = 4 \cdot 10^{-6}$, Figure S7) of environmental change alters our results with respect to the main scenarios in the main text ($\tau_c = 2 \cdot 10^{-6}$). We find that the patterns observed with slower evolutionary change are similar to those of faster environmental change, and vice versa. Results are thus determined by the relative timescales of evolution to the environment. In addition, we observe that changing one of the timescales at which evolution or environmental change acts mainly affects the effect size of extinction delay but not the pattern of evolutionary rescue and murder.

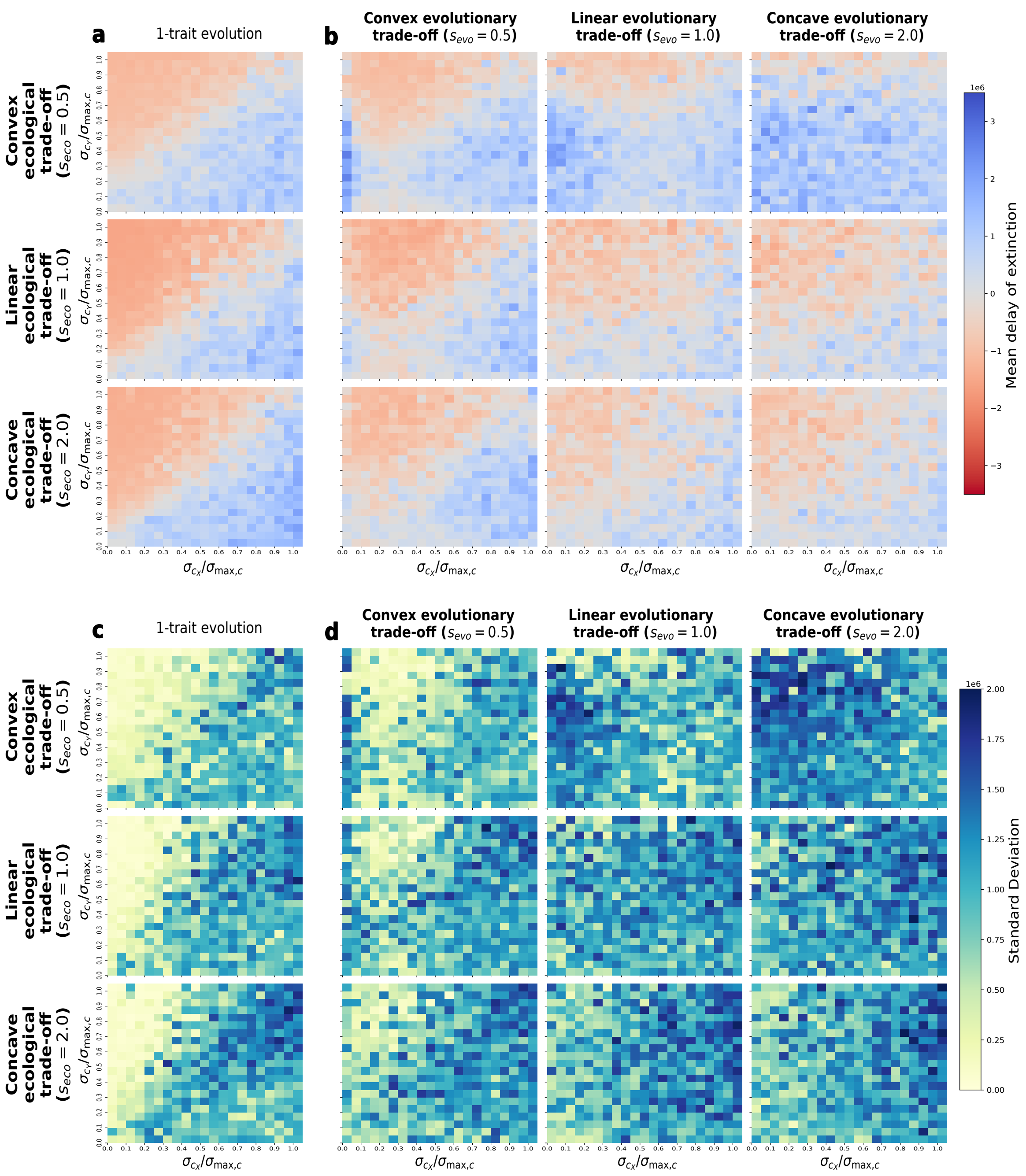

Figure S4: Effects of slower evolution on the eco-evolutionary dynamics. Extinction delay $\Delta t_X$ (i.e., difference in extinction time between evolutionary and ecological scenarios) for species $X$ in the 1-trait (a) and 2-trait (b) evolution scenario, and variability thereof over 10 replicates (c, d), with slower evolution ($\mu_X = \mu_Y = 5 \cdot 10^{-5}$ than in the main text. All other metrics are as for Figure 4 of the main text. In (a) and (c), given that only one trait evolves, no evolutionary trade-off exists, so only one heatmap is shown for each ecological trade-off shape. Colors represent (a, b) mean or (c, d) standard deviation of $\Delta t_X$ across 10 replicated simulation runs. Positive $\Delta t_X$ values (blue colors in a, b) indicate that evolution delays the extinction of species $X$, whereas negative values (red colors in a, b) indicate that evolution accelerates extinction.

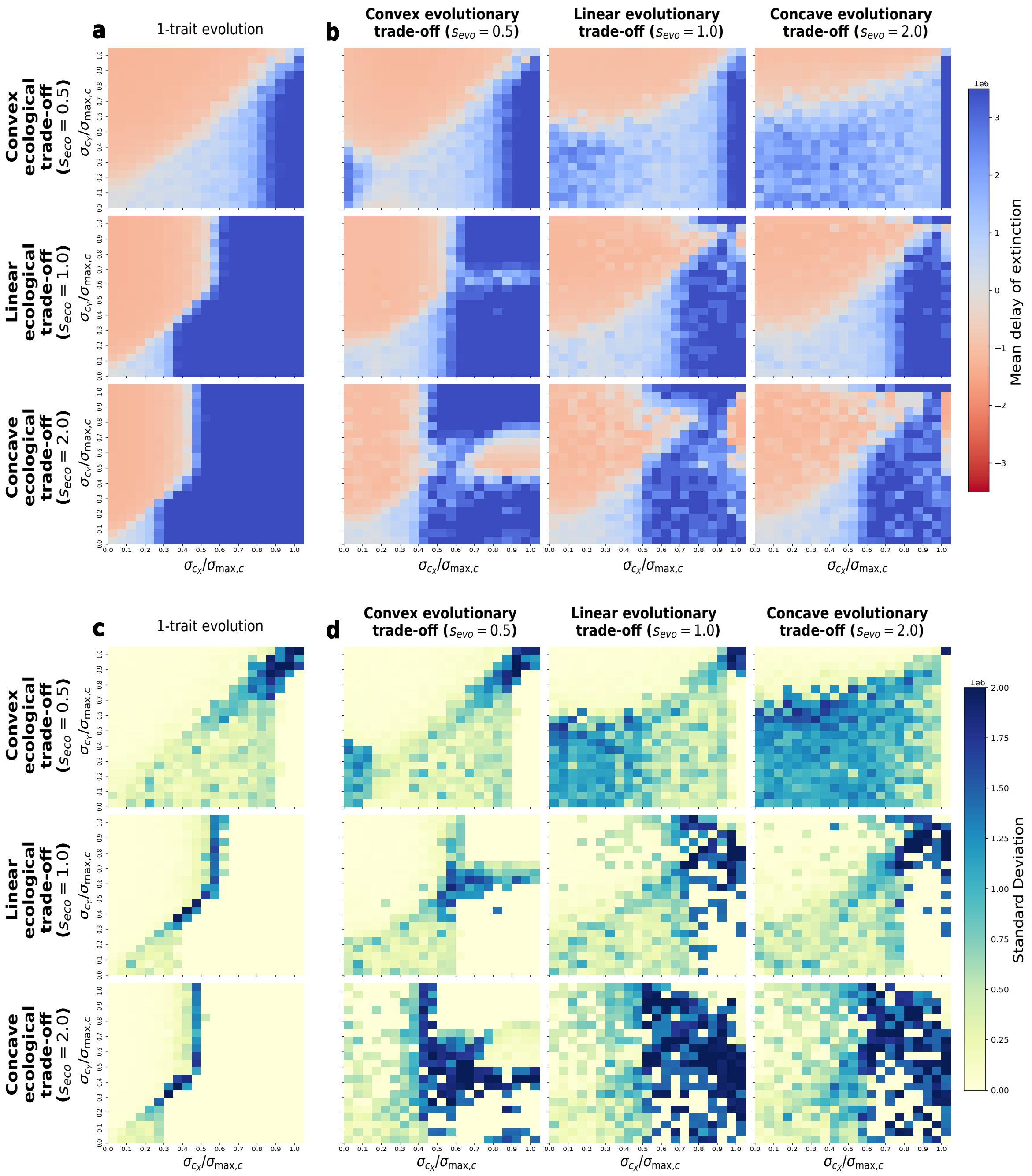

Figure S5: Effects of faster evolution on the eco-evolutionary dynamics. Extinction delay $\Delta t_X$ (i.e., difference in extinction time between evolutionary and ecological scenarios) for species $X$ in the 1-trait (a) and 2-trait (b) evolution scenario, and variability thereof over 10 replicates (c, d), with faster evolution ($\mu_X = \mu_Y = 1.5 \cdot 10^{-3}$) than in the main text. All other metrics are as for Figure 4 of the main text. In (a) and (c), given that only one trait evolves, no evolutionary trade-off exists, so only one heatmap is shown for each ecological trade-off shape. Colors represent (a, b) mean or (c, d) standard deviation of $\Delta t_X$ across 10 replicated simulation runs. Positive $\Delta t_X$ values (blue colors in a, b) indicate that evolution delays the extinction of species $X$, whereas negative values (red colors in a, b) indicate that evolution accelerates extinction.

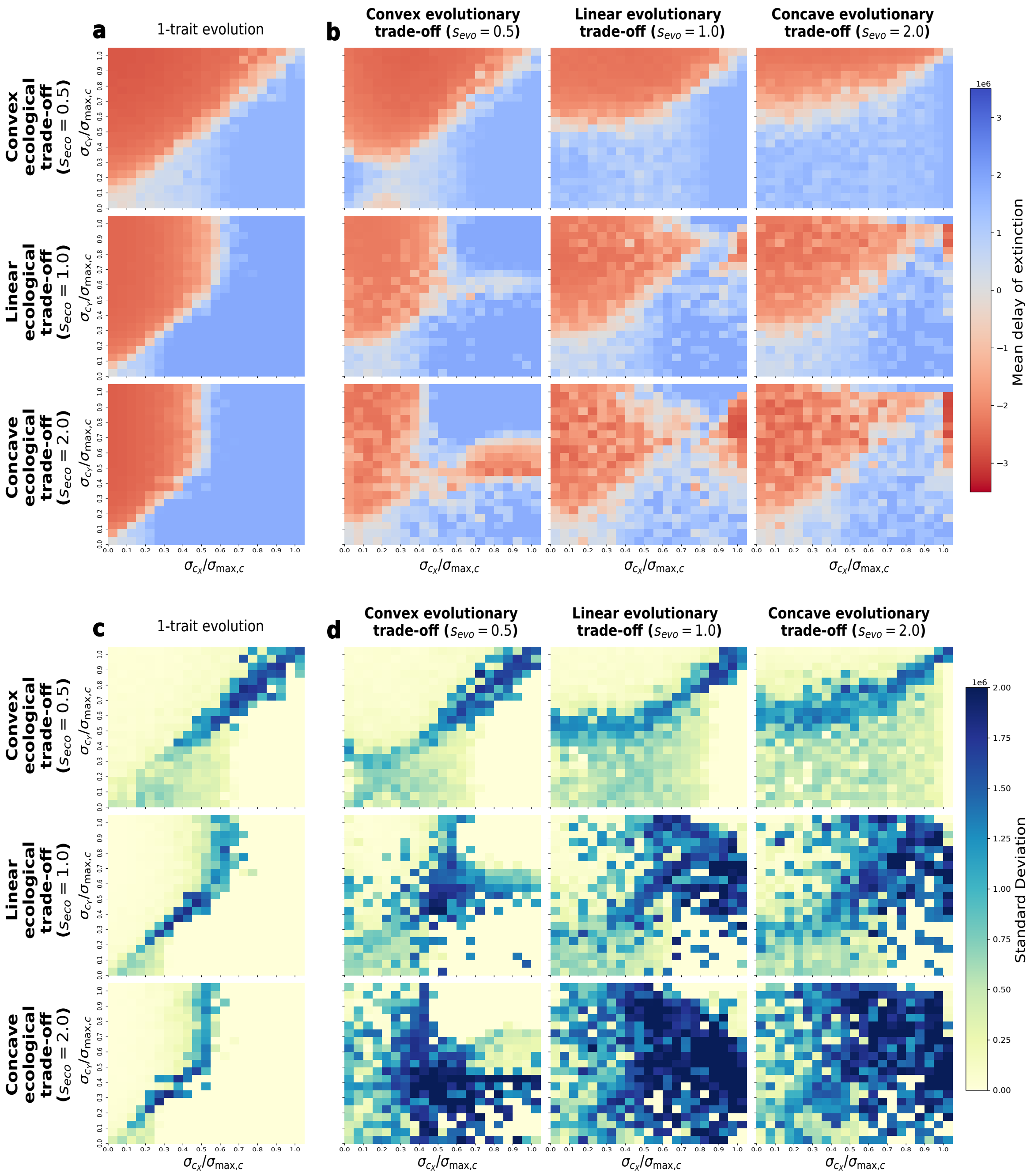

Figure S6: Effects of slower environmental change on the eco-evolutionary dynamics. Extinction delay $\Delta t_X$ (i.e., difference in extinction time between evolutionary and ecological scenarios) for species $X$ in the 1-trait (a) and 2-trait (b) evolution scenario, and variability thereof over 10 replicates (c, d), with slower environmental change ($\tau_c = 1 \cdot 10^{-6}$) than in the main text. All other metrics are as for Figure 4 of the main text. In (a) and (c), given that only one trait evolves, no evolutionary trade-off exists, so only one heatmap is shown for each ecological trade-off shape. Colors represent (a, b) mean or (c, d) standard deviation of $\Delta t_X$ across 10 replicated simulation runs. Positive $\Delta t_X$ values (blue colors in a, b) indicate that evolution delays the extinction of species $X$, whereas negative values (red colors in a, b) indicate that evolution accelerates extinction.

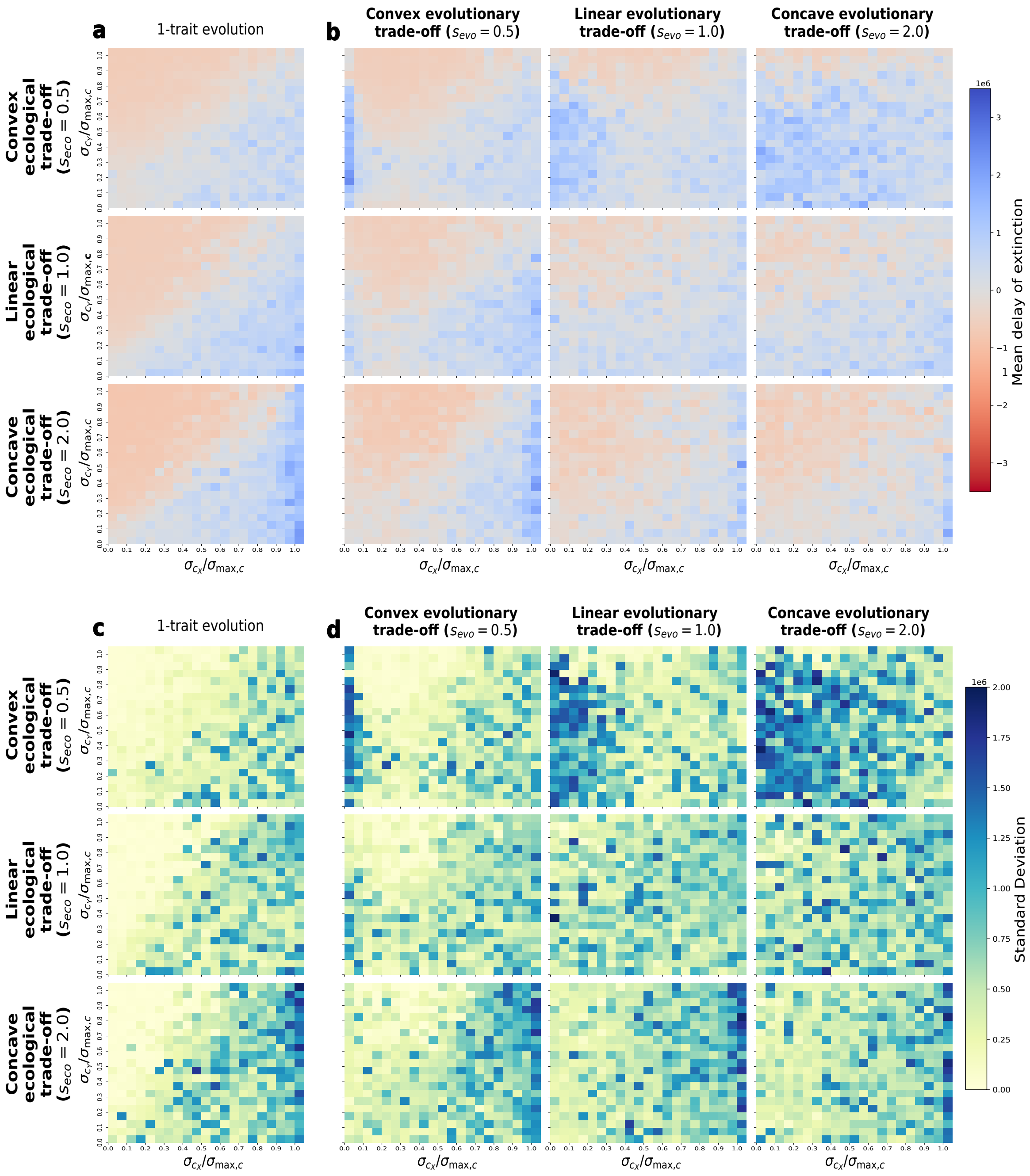

Figure S7: Effects of faster environmental change on the eco-evolutionary dynamics. Extinction delay $\Delta t_X$ (i.e., difference in extinction time between evolutionary and ecological scenarios) for species $X$ in the 1-trait (a) and 2-trait (b) evolution scenario, and variability thereof over 10 replicates (c, d), with faster environmental change ($\tau_c = 4 \cdot 10^{-6}$) than in the main text. All other metrics are as for Figure 4 of the main text. In (a) and (c), given that only one trait evolves, no evolutionary trade-off exists, so only one heatmap is shown for each ecological trade-off shape. Colors represent (a, b) mean or (c, d) standard deviation of $\Delta t_X$ across 10 replicated simulation runs. Positive $\Delta t_X$ values (blue colors in a, b) indicate that evolution delays the extinction of species $X$, whereas negative values (red colors in a, b) indicate that evolution accelerates extinction.

#### S9 Stable coexistence for the initial types

Here, we outline the derivations for the conditions under which two initial types coexist stably if there is no evolution. These derivations aim to establish the conditions that justify the chosen initial values of $d_X, d_Y, b_{XY}$ and $b_{YX}$, ensuring that the species are initialized in stable coexistence so that any differences in ecological outcomes are due to evolution. Initially, only one type of each species is present, so that the governing equations are those of the following Lotka-Volterra model adapted to our system (see Equation (1) in the main text):

$$\begin{cases} \dfrac{\mathrm{d}x}{\mathrm{d}t} = r_X x \left( w_X - a_X(x + b_{XY} y) \right), \\ \dfrac{\mathrm{d}y}{\mathrm{d}t} = r_Y y \left( w_Y - a_Y(y + b_{YX} y) \right). \end{cases}$$

The dynamics of these equations are well known and included in many textbooks on the subject matter, such as Kot (2001). In particular, the principle that "mutual invasion implies coexistence" holds (Chesson & Kuang 2008). For convenience of the reader and for completeness, we present the most important calculations here.

The system has up to four biologically relevant equilibrium points, i.e., all points $(x, y)$ with $x, y \geq 0$, where $\mathrm{d}x/\mathrm{d}t = 0 = \mathrm{d}y/\mathrm{d}t$. They are given by the trivial equilibrium $(0, 0)$ where both species are extinct, the two semi-trivial equilibria

$$E_X = (\widehat{x}, 0) = (d_X w_X, 0), \quad \text{and} \quad E_Y = (0, \widehat{y}) = (0, d_Y w_Y),$$

where only one of the two species is present, and the coexistence equilibrium $(x^*, y^*)$ with

$$x^* = \frac{d_X w_X - b_{XY} d_Y w_Y}{1 - b_{XY} b_{YX}}, \quad \text{and} \quad y^* = \frac{d_Y w_Y - b_{YX} d_X w_X}{1 - b_{XY} b_{YX}}. \tag{S9.1}$$

While the semi-trivial equilibria are always nonnegative, the coexistence equilibrium is positive only if numerators and the denominator are either all positive or all negative, but not in the mixed case.

To investigate the stability of each equilibrium, we calculate the Jacobian matrix

$$J(x,y) = \begin{pmatrix} r_X(w_X - a_X\,(2x + b_{XY}y)) & -r_X a_X b_{XY} x \\ -r_Y a_Y b_{YX} y & r_Y(w_Y - a_Y\,(2y + b_{YX}x)) \end{pmatrix}.$$

At the trivial equilibrium, the matrix becomes

$$J(0,0) = \begin{pmatrix} r_X w_X & 0 \\ 0 & r_Y w_Y \end{pmatrix}.$$

Since the matrix is diagonal, the diagonal entries are the eigenvalues. Since both are positive, the equilibrium is unstable, meaning each species can grow from small densities.

At $E_X$, the Jacobian matrix becomes

$$J(\widehat{x},0) = \begin{pmatrix} -r_X w_X & -r_X w_X b_{XY} \\ 0 & r_Y\,(w_Y - a_Y b_{YX} d_X w_X) \end{pmatrix}.$$

Since the matrix is upper triangular, the diagonal entries are again the eigenvalues. The first is negative, indicating that $E_X$ is stable with respect to perturbations in the $x$-direction. The second gives the per-capita growth rate of species $Y$ at $E_X$. Species $Y$ can grow from low density and therefore *invade* $X$ precisely when this quantity is positive. Since $r_Y$ is positive, species $Y$ can invade $X$ when

$$d_Y w_Y - b_{YX} d_X w_X > 0. \tag{S9.2}$$

The case of $E_Y$ is analogous, so that species $X$ can invade species $Y$ if

$$d_X w_X - b_{XY} d_Y w_Y > 0. \tag{S9.3}$$

To find the stability conditions at the coexistence point, we use the Routh-Hurwitz conditions that the equilibrium is stable if and only if $\mathrm{tr}(J(x^*, y^*)) < 0$ and $\det(J(x^*, y^*)) > 0$. We rearrange the expressions of the coexistence equilibrium (S9.1) to get

$$w_X = a_X(x^* + b_{XY}y^*) \quad \text{and} \quad w_Y = a_Y(y^* + b_{YX}x^*). \tag{S9.4}$$

When we substitute these expressions into the Jacobian matrix, we obtain

$$J(x^*, y^*) = \begin{pmatrix} -r_X x^* a_X & -r_X b_{XY} x^* a_X \\ -r_Y b_{YX} y^* a_Y & -r_Y y^* a_Y \end{pmatrix}. \tag{S9.5}$$

We calculate the trace and the determinant as

$$\mathrm{tr}(J(x^*, y^*)) = -r_X x^* a_X - r_Y y^* a_Y, \qquad \det(J(x^*, y^*)) = r_X r_Y x^* y^* a_X a_Y (1 - b_{XY} b_{YX}).$$

To evaluate the stability of the coexistence state, we note that since $x^*, y^* > 0$, the condition $\mathrm{tr} J(x^*, y^*) < 0$ is automatically satisfied. The condition $\det J(x^*, y^*) > 0$ is satisfied if and only if $1 - b_{XY}b_{YX} > 0$. The latter implies that, necessarily,

$$d_X w_X - b_{XY} d_Y w_Y > 0, \quad \text{and} \quad d_Y w_Y - b_{YX} d_X w_X > 0, \tag{S9.6}$$

because of the expressions in (S9.1). Hence, the positive coexistence point is stable if and only if $1 - b_{XY}b_{YX} > 0$.

Comparing the inequalities (S9.6) with those in (S9.3) and (S9.2), we see that if there is a stable positive coexistence state, then we have mutual invasibility between species $X$ and $Y$. It turns out that the reverse of this statement also holds: Mutual invasibility implies the existence of a stable coexistence equilibrium. To see this, we start with the expressions (S9.3) and (S9.2). From (S9.2), we have $d_Y w_Y > b_{YX} d_X w_X$. Multiplying (S9.3) by $b_{YX} > 0$ and rearranging, we get $b_{YX}d_Xw_X > b_{XY}b_{YX}d_Yw_Y$. Hence, we have the chain of inequalities

$$d_Yw_Y > b_{YX}d_Xw_X > b_{XY}b_{YX}d_Yw_Y.$$

Comparing the first with the last term, we find that necessarily

$$1 - b_{XY}b_{YX} > 0.$$

Hence, the coexistence state is positive and, by the above stability conditions, also stable.

We can also rewrite the mutual invasion conditions (S9.3) and (S9.2) as

$$b_{YX} < \frac{d_Yw_Y}{d_Xw_X} \quad \text{and} \quad \frac{d_Yw_Y}{d_Xw_X} < \frac{1}{b_{XY}},$$

which can be combined into the desired condition

$$b_{YX} < \frac{d_Yw_Y}{d_Xw_X} < \frac{1}{b_{XY}}.$$

If we choose our species $X$ and $Y$ to be initially identical, we have $d_X = d_Y$, $w_X = w_Y$, and $b_{XY} = b_{YX}$, so that this chain of inequalities is satisfied whenever $0 < b_{XY} < 1$.
